# ΔNp63 drives dysplastic alveolar remodeling and restricts epithelial plasticity upon severe lung injury

**DOI:** 10.1101/2022.02.23.481695

**Authors:** Aaron I. Weiner, Gan Zhao, Hanna M. Zayas, Nicolas P. Holcomb, Stephanie Adams-Tzivelekidis, Joanna Wong, Maria E. Gentile, Gargi Palashikar, Kwaku Quansah, Andrew E. Vaughan

## Abstract

Unlike many mammalian vital organs, the lung exhibits a robust, multifaceted regenerative response to severe injuries such as influenza infection, which primarily targets epithelial cells in the airways and alveoli. Quiescent lung-resident epithelial progenitors proliferate, migrate, and differentiate following lung injury, participating in two distinct reparative pathways: functionally beneficial regeneration and dysplastic tissue remodeling. Intrapulmonary airway-resident basal-like p63^+^ progenitors are one such progenitor cell type that migrates from the airways to form ectopic bronchiolar tissue in the alveoli, generating honeycomb-like cysts that fail to resolve after injury. Though this phenomenon is now well described, the cell-autonomous signals that drive dysplastic alveolar remodeling remain uncertain, a question made especially salient by observations that p63^+^ progenitors also expand dramatically upon diffuse alveolar damage in humans resulting from a variety of insults including SARS-CoV-2-induced ARDS. Here we show that the master basal cell transcription factor ΔNp63 is required for the immense migratory capacity of intrapulmonary p63^+^ progenitors and consequently for the dysplastic repair pathway manifest by these cells. We further demonstrate that ΔNp63 restricts the fate plasticity of intrapulmonary p63^+^ progenitors by regulating their epigenetic landscape, and that loss of ΔNp63 alters the deposition of active and repressive histone modifications at key differentiation gene loci, allowing ΔNp63^KO^ progenitors to proceed towards airway or alveolar differentiation depending on their surrounding environment. These insights into the regulatory mechanisms of dysplastic repair and intrapulmonary p63^+^ progenitor fate choice highlight potential therapeutic targets to promote more effective alveolar regeneration following severe lung injuries.

## Introduction

Lung alveolar epithelial repair involves a complex interplay of multiple progenitor cell types from distinct anatomical compartments, primarily the airways and alveoli, synergizing to ultimately re-epithelialize the alveolar barrier after pulmonary injury. Severe respiratory infections and epithelial insults primarily target the epithelial cells of the gas-exchanging alveoli, alveolar type two (AT2) and type-one (AT1) cells. Two broad regenerative pathways are subsequently simultaneously undertaken to repair the alveoli: functionally beneficial regeneration of the alveolar epithelium and dysplastic alveolar remodeling that follows severe lung injuries such as H1N1 influenza. AT2s and club-like progenitor cells residing near the bronchioalveolar duct junction (distal club/BASCs) are the primary players in functionally beneficial (i.e. “euplastic”) regeneration, during which they proliferate and differentiate into AT1s (Barkauskas et al., 2013; Evans et al., 1973; Kathiriya et al., 2020; Kim et al., 2005; Liu et al., 2019), restoring both the surfactant production and gas-exchanging alveolar epithelium. Following severe injuries in which AT2s are entirely ablated in large regions (diffuse alveolar damage, DAD), dysplastic repair employs a rare population of basal cell-like p63^+^ progenitor cells present in the intrapulmonary airways, which undergo widespread migration into the alveolar compartment and generate ectopic airway-like epithelium comprised of p63^+^ Krt5^+^ basal cells (Vaughan et al., 2015; Xi et al., 2017; Zuo et al., 2015) as well as mature airway epithelial cells, including Scgb1a1^+^ Scgb3a2^+^ club cells (Weiner et al., 2019) and Dclk1^+^ Trpm5^+^ tuft cells (Rane et al., 2019). This dysplastic bronchiolization persists indefinitely as honeycomb-like cysts (Kanegai et al., 2016; Vaughan et al., 2015) that do not appear to contribute to lung function, instead inhabiting alveolar space where functionally beneficial regeneration might otherwise occur. This maladaptive repair by intrapulmonary p63^+^ progenitors is similar to alveolar bronchiolization seen in humans with interstitial lung disease (ILD) (Seibold et al., 2013; Smirnova et al., 2016; Vaughan et al., 2015), acute respiratory distress syndrome (ARDS) and DAD (Taylor et al., 2018), and COVID-19 (Fernanda de Mello Costa et al., 2020; Melms et al., 2021; Zhao et al., 2020), in which dysplastic bronchiolization forms ectopically in the alveoli. The loss of alveolar epithelial populations and the failure of ectopic bronchiolized tissue to resolve over time suggest that while it probably aids in barrier restoration in the short term, this dysplastic repair response is ultimately maladaptive, likely contributing to the long-term compromise of pulmonary function seen in some survivors of severe pulmonary injury (Koppe et al., 2016; Liu et al., 2015).

While the vast majority of intrapulmonary p63^+^ progenitors contribute to dysplastic alveolar remodeling and either remain as p63^+^ Krt5^+^ basal cells or differentiate into mature airway lineages within the alveoli, lineage tracing experiments have demonstrated the exceedingly rare contribution of post-injury intrapulmonary p63^+^ progenitors to surfactant protein C (SPC)^+^ AT2s in the alveoli (Vaughan et al., 2015; Yang et al., 2018), raising the possibility that intrapulmonary p63^+^ progenitors might exhibit some inherent plasticity. The effects of paracrine signaling pathways such as Wnt, Notch, and Fgf on dysplastic remodeling have been the focus of much recent attention (Quantius et al., 2016; Vaughan et al., 2015; Xi et al., 2017; Yuan et al., 2019), though little is known about the cell-intrinsic regulators necessary to mount dysplastic expansion or resolve fate outcomes *in vivo*. Given their long-term persistence after injury and likely contribution to jeopardized lung function, capitalizing on their limited but demonstrable plasticity and altering the fate of injury-activated intrapulmonary p63^+^ progenitors is an attractive avenue to promote functionally beneficial, euplastic regeneration.

Intralobular p63^+^ progenitors are the only cells in the lung proper that express the master basal cell identity transcription factor p63, which is highly expressed in basal stem cells of proliferative epithelial tissues such as the skin (Pellegrini et al., 2001), prostate (Signoretti et al., 2005; Signoretti et al., 2000), mammary gland (Barbareschi et al., 2001; Van Keymeulen et al., 2011), and trachea (Rock et al., 2009). ΔNp63 is the predominantly expressed *Trp63* isoform in embryonic (Laurikkala et al., 2006) and adult (Yang et al., 1998) epithelial cells, including intrapulmonary p63^+^ progenitors (Vaughan et al., 2015), and is a crucial driver of basal cell identity (Romano et al., 2012), proliferation (Truong et al., 2006; Yang et al., 1999), migration (Carroll et al., 2006; Giacobbe et al., 2016; Gu et al., 2008), and differentiation (Haas et al., 2019; Kouwenhoven et al., 2015; Romano et al., 2012; Signoretti et al., 2005; Truong et al., 2006) in both homeostatic and regenerative contexts. ΔNp63 is necessary to maintain basal cell identity and function; without ΔNp63, these cells downregulate basal markers, exit the cell cycle, and undergo a program of precocious differentiation into more committed epithelial cells of their resident tissue (Haas et al., 2019; Romano et al., 2012; Truong et al., 2006; Yang et al., 2018). While ΔNp63 exterts its control over basal cell identity directly by transcriptionally regulating expression of basal cell markers and functional components such as basement membrane-anchoring adhesion molecules (Carroll et al., 2006; Ferone et al., 2013; Ihrie et al., 2005; Kurata et al., 2004; Laurikkala et al., 2006; Romano et al., 2009), recent studies propose an epigenetic role for ΔNp63 as well, in which ΔNp63 cooperates with higher order chromatin remodelers and histone modifiers to rearrange the 3D topology and chromatin accessibility of the genome (Fan et al., 2018; Fessing et al., 2011; Mardaryev et al., 2014; Qu et al., 2019; Sethi et al., 2014), bookmark tissue-specific enhancer regions during differentiation (Kouwenhoven et al., 2015; Lin-Shiao et al., 2019), and acts as a pionner-like factor by binding and opening inaccessible chromatin (Fan et al., 2018; Santos-Pereira et al., 2019; Yu et al., 2021). With its multi-tiered influence over the fundamental basal cell program and critical role in virtually all epithelial tissue contexts in which its been studied, ΔNp63 serves as a promising target to modulate basal cell fate choice in regions of dysplastic remodeling within injured lungs.

In this study, we define the foundational role of ΔNp63 in the activation and maintenance of intrapulmonary p63^+^ progenitors and demonstrate that loss of ΔNp63 blunts the dysplastic remodeling program, restricting the migratory capacity of intrapulmonary p63^+^ progenitors and immobilizing them in the airways, allowing for a more substantive contribution from other (p63^negative^) airway progenitors to AT2 reconstitution. Further, we provide evidence that by alleviating the basal transcriptional module conferred by ΔNp63, intrapulmonary p63^+^ progenitors can be “pushed towards” appropriate alveolar epithelial cell fates. We confirm the near-permanent persistence of dysplasia in injured murine lungs and demonstrate elevated levels of ΔNp63 in injured mouse and human basal cells, including in the context of COVID-19. ChIP-qPCR analysis of H3K27me3 and H3K27ac deposition at key lung epithelial differentiation genes reveals rewiring of the epigenetic landscape with ΔNp63 deletion, potentiating multiple differentiation trajectories. Indeed, ΔNp63^KO^ intrapulmonary p63^+^ progenitors assumed a club cell-like state, which could be guided towards either differentiation into airway cell lineages or toward an AT2 lineage depending on *in vitro* conditions. Taken together, our findings elucidate the role of ΔNp63 in intrapulmonary p63^+^ progenitors and the dysplastic remodeling program, demonstrating that the basal cell program can be reversed and the plasticity of intrapulmonary p63^+^ progenitors unlocked, making ΔNp63 an attractive therapeutic candidate for individuals with severe pulmonary injuries.

## Results

### Dysplastic alveolar remodeling persists indefinitely in mice

Dysplastic remodeling of the alveolar epithelial compartment occurs in response to severe pulmonary injuries in mice and humans and is thought to persist as an “epithelial scar” long after the initial injury event (Kanegai et al., 2016; Vaughan et al., 2015). We confirmed that ΔNp63^+^ Krt5^+^ dysplastic pods remain in 100% (n = 3/3) of wild-type mice after injury with H1N1 influenza A virus strain PR8 (Gerber et al., 1955; Loosli et al., 1975) one year (nearly half of the lifespan of adult mice) post-infection (Fig. 1a-a’). We found proliferating Krt5^+^ Ki67^+^ dysplastic cells in all surveyed mice (n = 3/3) (Fig. 1b-d’), confirming the self-renewal of these cells to maintain their foothold in injured alveolar regions long after the initial injury. The endurance of epithelial dysplasia was accompanied by a significant reduction oxygen saturation at one-year post-infection compared to age-matched uninjured mice as assessed by pulse oximetry (Fig. 1e), reflecting the chronic physiological consequences of severe alveolar injury.

**Figure 1.**
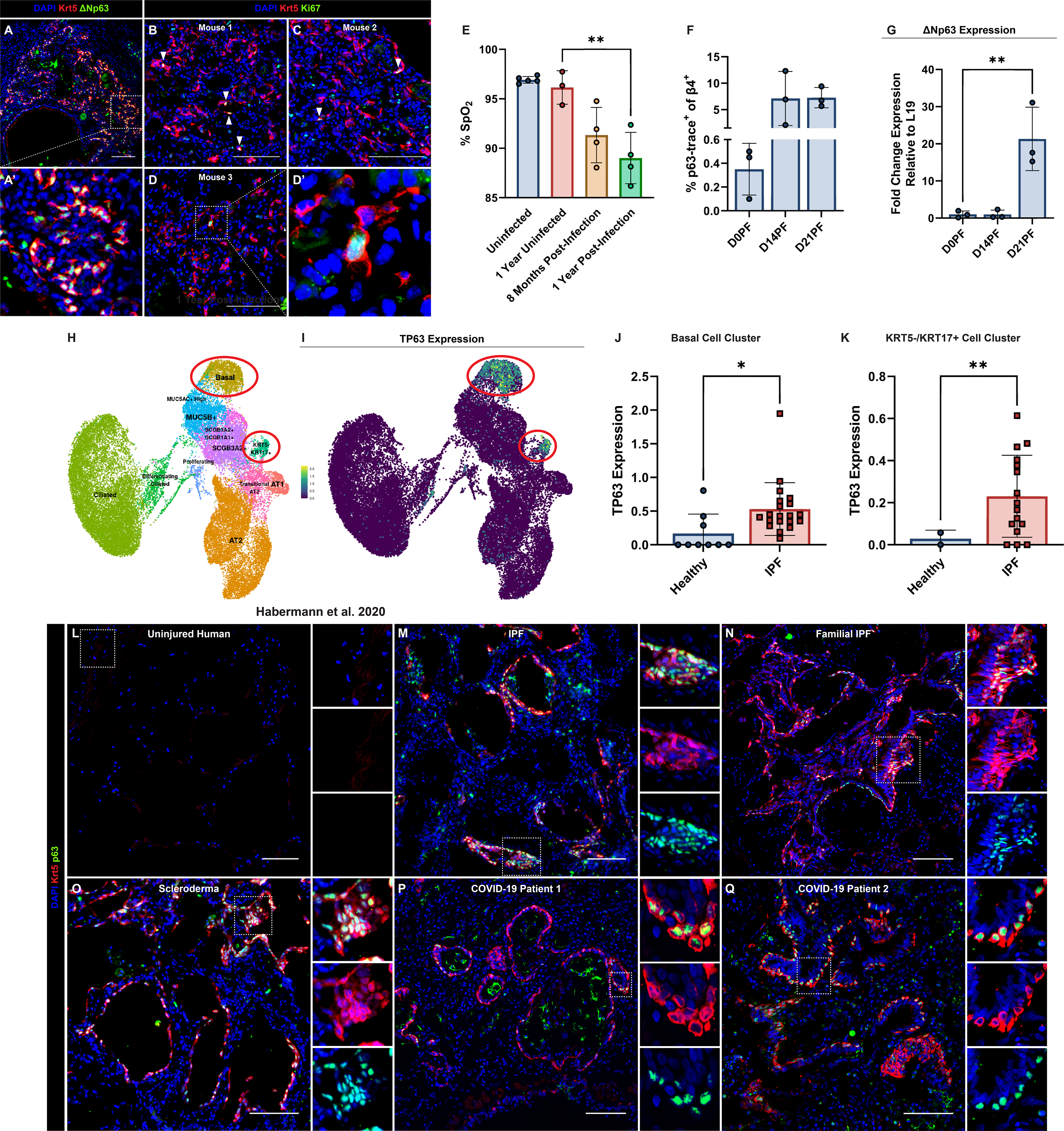
Dysplastic alveolar remodeling persists long-term in injured mice and humans and is correlated with ΔNp63 expression. (a) ΔNp63^+^ Krt5^+^ dysplastic remodeling persists in murine alveolar regions one-year post-flu. Scale bar = 100um. (a’) Inset of outlined region in (a). (b-d) Representative images of Krt5^+^ Ki67^+^ proliferating dysplastic cells (white arrows) from three mice (100% of mice surveyed). Scale bar = 100um. (d’) 63x magnification inset image of outlined region in (d). (e) Pulse oximetry readings were taken on uninjured two-to-three-month-old mice, uninjured one year old mice, injured mice at eight months post-infection, and the same mice at one-year post- infection. Significance based on ordinary one-way ANOVA. (f) Flow cytometry quantification during injury time course of % p63-trace^+^ as a fraction of live CD45^-^ EpCAM^+^ β4^+^ cells. (g) ΔNp63 expression increases in sorted p63-trace^+^ cells with injury progression. Significance based on ordinary one-way ANOVA. (h) Human healthy and IPF epithelial cell UMAP from IPF Cell Atlas (Habermann et al., 2020) with basal and KRT5-/KRT17+ clusters highlighted. (i) TP63 expression is restricted to basal and KRT5-/KRT17+ clusters in cells from both healthy and IPF patients. From IPF Cell Atlas (Habermann et al., 2020). (j, k) TP63 expression is significantly upregulated in IPF patient epithelial cells compared to healthy epithelial cells from both the (j) basal and (k) KRT5-/KRT17+ clusters. Significance for both (j) and (k) based on Welch’s t-test. (l-q) Representative immunofluorescence staining for Krt5 and p63 in alveolar regions of (l) uninjured human, (m) IPF patient, (n) familial IPF patient, (o) scleroderma patient, and (p, q) two COVID-19 patient lungs. Krt5^+^ p63^+^ dysplastic pods are only found in injured alveoli. Insets are of outlined regions. Scale bars for (l-q) = 100um. Data represented as mean ± standard deviation and *p < 0.05, **p < 0.01, ***p < 0.001, ****p < 0.0001 for all statistics.

### Intrapulmonary p63^+^ progenitors are not directly infected by influenza A virus

Intrapulmonary p63^+^ cells are very rare in uninjured lungs but expand dramatically upon injury, suggesting they may be resistant to influenza infection. To determine if intrapulmonary p63^+^ progenitors are initially infected, we utilized an engineered PR8 strain that expresses a copy of Cre recombinase (IAV-Cre) (Hamilton et al., 2016; Heaton et al., 2014). When administered to transgenic lineage reporter mice, cells that are actively infected by an IAV-Cre virion will express Cre recombinase, allowing for lineage tracing of infected cells (Supplementary Fig. 1a). At day 28 after IAV-Cre infection of ai14tdTomato (RFP) mice, we observed large IAV-trace^+^ regions of the alveoli comprised mostly of SPC^+^ AT2s and RAGE^+^ AT1s (Supplementary Fig. 1b), likely arising from infected AT2s and distal club cells/BASCs but also potentially direct infection of AT1s. The vast majority of Krt5^+^ dysplastic cells in IAV-Cre-infected mice were untraced (Supplementary Fig. 1c), though rare IAV-trace^+^ Krt5^+^ regions were observed (Supplementary Fig. 1d), suggesting that intrapulmonary p63^+^ progenitors are not the primary target of PR8 but that direct infection is possible. Quantification of the fraction of IAV-traced cells bearing lineage markers in the airways and alveoli supported this conclusion, as Scgb3a2^+^ club cells in the airways and SPC^+^ AT2s in the alveoli were the most abundantly labeled epithelial cells in their respective compartments while Krt5^+^ cells constituted a small fraction (∼2% of IAV-traced cells in the airways and ∼6% of IAV-traced cells in the alveoli) of labelled cells in these regions (Supplementary Fig. 1e, f). Taken together, this indicates that intrapulmonary p63^+^ progenitors largely escape initial infection, but rare cells that are infected are able to clear virus and survive indefinitely.

### p63 expression is increased in injured mouse and human basal cells

We next sought to uncover cell-intrinsic molecular drivers of the expansion and maintenance of dysplastic remodeling after pulmonary injury. The basal cell transcription factor p63, specifically its highly transcriptionally active ΔN isoform, is well-known for its role in maintaining basal cell identity and promoting many characteristics common to basal cells during wound healing, including basal cell proliferation, migration, and differentiation. Since intrapulmonary p63^+^ progenitors are the only cells in the distal lung that express ΔNp63, we chose to investigate the role of ΔNp63 in intrapulmonary p63^+^ progenitors and their contribution to dysplasia. Using the p63CreERT2 mouse to lineage trace and isolate intrapulmonary p63^+^ progenitors during injury, we found that intrapulmonary p63^+^ progenitors multiply almost 50-fold from the beginning of injury to the end of viral clearance and injury resolution, ultimately encompassing almost 10% of cells marked by airway epithelial integrin β4 (β4) (Fig. 1f) though this is likely an underestimate due to the haploinsufficiency previously noted in p63^CreERT2^ mice (Yang et al., 2018). Further, ΔNp63 expression increased dramatically on a “per-cell” basis as intrapulmonary p63^+^ progenitors expanded ectopically into the alveoli, reaching its peak expression by day 21 post-flu (Fig. 1g). To determine if p63 expression similarly increased in injured human basal cells, we re-analyzed single-cell RNA sequencing datasets from healthy human and human IPF patient lung tissue samples (Habermann et al., 2020). p63 expression is restricted to two epithelial populations within this dataset: basal cells and a population identified as Krt5-/Krt17+ cells (Fig. 1h, i), a basal-like population that is largely restricted to diseased patient samples and that is associated with “transitional” AT2s in mice and humans and transdifferentiation of AT2s into basaloid cells in humans (Choi et al., 2020; Kathiriya et al., 2022; Kobayashi et al., 2020; Strunz et al., 2020). We confirmed that p63 expression is significantly elevated in human basal cells (Fig. 1j) and Krt5-/Krt17+ cells (Fig. 1k) from IPF patients compared to healthy donors, indicating that p63^+^ progenitors and p63 expression itself are dynamically regulated in both mouse and human lungs upon injury. We also observed strong p63 staining in the Krt5^+^ cells that line the dysplastic epithelial cysts in the alveoli of human patients with a variety of pulmonary injuries and diseases, including IPF (Fig. 1m), familial IPF (Fig. 1n), scleroderma (Fig. 1o), and COVID-19 (Fig. 1p, q), but not in the alveolar regions of healthy patients (Fig.1l), further supporting the potential relevance of p63 in human lung disease.

### Broad airway epithelial ΔNp63 deletion attenuates dysplastic alveolar remodeling

Since the dysplastic cell-of-origin resides in the intrapulmonary airway epithelium (Ray et al., 2016; Xi et al., 2017), we asked whether broad ΔNp63 deletion in the airway epithelium could attenuate alveolar Krt5^+^ dysplasia following influenza injury *in vivo*. Utilizing Sox2^CreERT2^ mice bearing RFP and ΔNp63^flox^ alleles, we simultaneously lineage traced and deleted ΔNp63 in the airway epithelium (Arnold et al., 2011; Chakravarti et al., 2014; Madisen et al., 2010). Sox2^CreERT2^; RFP (Sox2^ΔNp63WT^) and Sox2^CreERT2^; RFP; ΔNp63^flox/flox^ (Sox2^ΔNp63KO^) mice were administered a single dose of intraperitoneal (IP) tamoxifen and infected with PR8 (Fig. 2a). While Sox2^ΔNp63WT^ mice had large expansions of Sox2- trace^+^ Krt5^+^ dysplasia consistent with their severe injury (Fig. 2b), Sox2^ΔNp63KO^ mice did not display any Krt5^+^ dysplasia (Fig. 2c), prompting us to examine the lineage contributions that Sox2^ΔNp63WT^ and Sox2^ΔNp63KO^ mice make to alveolar repair. Consistent with previous reports (Ray et al., 2016; Xi et al., 2017), influenza-injured Sox2^ΔNp63WT^ mice displayed many Sox2-trace^+^ SPC^+^ AT2s in the alveoli (accounting for ∼34% of Sox2-derived alveolar cells) (Fig. 2d, h) along with Sox2-trace^+^ Krt5^+^ bronchiolized dysplasia (accounting for ∼41% of Sox2-derived alveolar cells) (Fig. 2e, h) at day 21 post-flu, likely originating from injury-activated distal club/bronchioalveolar stem cells (BASCs) (Kathiriya et al., 2020; Liu et al., 2019; Rawlins et al., 2009) and intrapulmonary p63^+^ progenitors (Kumar et al., 2011; Vaughan et al., 2015; Xi et al., 2017), respectively. While regions of Sox2-trace^+^ SPC^+^ alveolar expansion persisted in Sox2^ΔNp63KO^ mice (Fig. 2f), Sox2-trace^+^ Krt5^+^ dysplastic alveolar regions were exceedingly rare (Fig. 2g), decreasing from ∼41% of Sox2-trace^+^ alveolar cells in Sox2^ΔNp63WT^ mice to ∼5% of Sox2-trace^+^ alveolar cells in Sox2^ΔNp63KO^ mice (Fig. 2h), indicating that ΔNp63 is necessary for airway epithelial cells to contribute to maladaptive alveolar remodeling, and that all Krt5^+^ dysplasia is derived from Sox2^+^ intrapulmonary p63^+^ progenitors.

**Figure 2.**
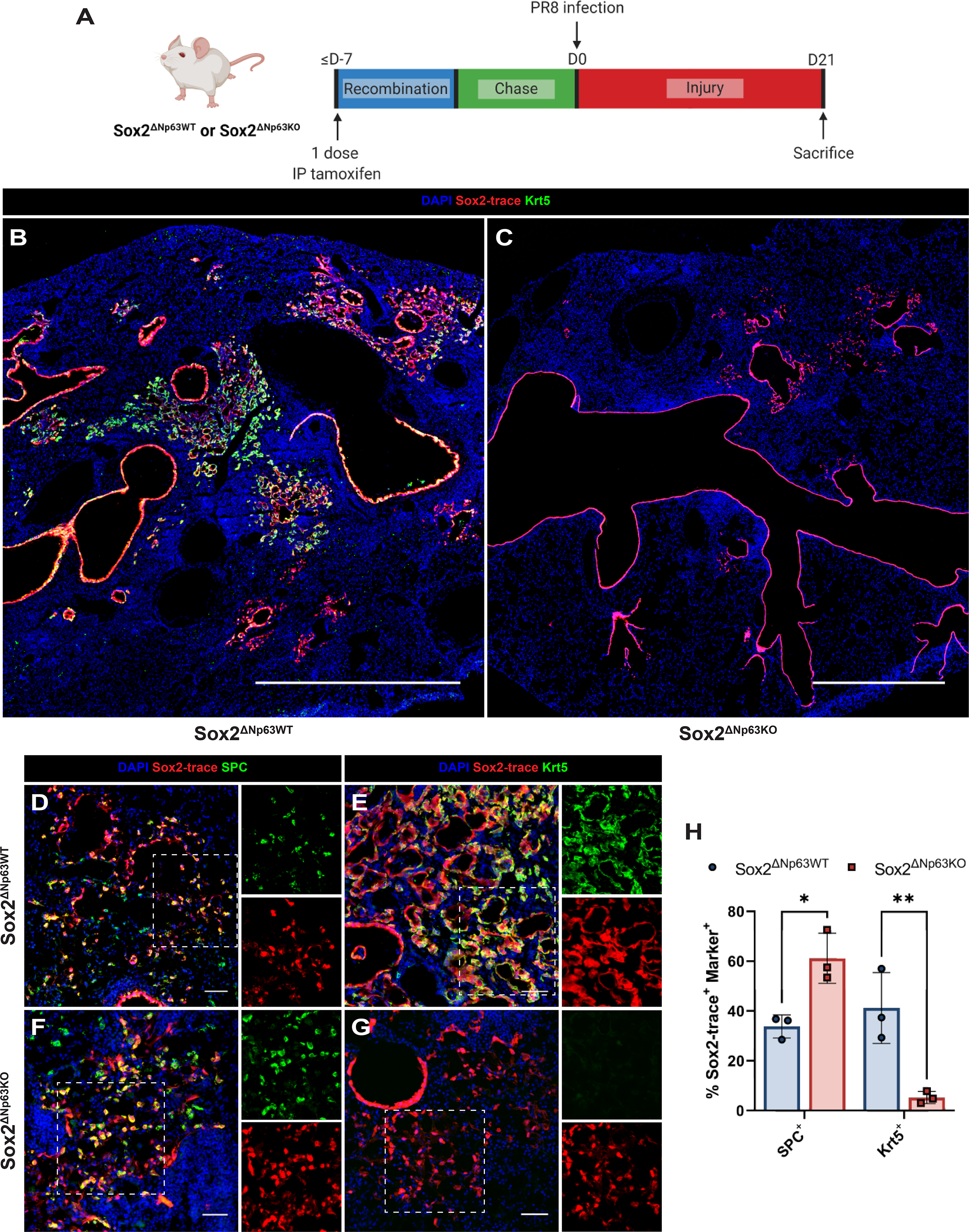
ΔNp63 is required for airway epithelial contribution to dysplastic remodeling *in vivo*. (a) Experimental timeline of Sox2-mediated ΔNp63 deletion. (b) 20x tilescan of a Sox2^ΔNp63WT^ lobe shows normal Sox2-trace^+^ contribution to alveolar repair, including Krt5^+^ dysplasia. (c) Sox2^ΔNp63KO^ lungs bear no Krt5^+^ dysplastic areas. Scale bar for (b) and (c) = 1mm. (d-g) Representative images of alveolar Sox2-trace overlap with lineage markers. Sox2^ΔNp63WT^ mice exhibit expansion of both (d) Sox2- trace^+^ SPC^+^ AT2s and (e) Sox2-trace^+^ Krt5^+^ dysplastic cells, whereas Sox2^ΔNp63KO^ mice have only (f) Sox2-trace^+^ SPC^+^ AT2s but lack (g) Sox2-trace^+^ Krt5^+^ dysplastic cells. Insets in (d-g) are of outlined regions. Scale bars for (d-g) = 50um. (h) Quantification of Sox2-trace contribution to SPC^+^ AT2s and Krt5^+^ dysplasia. Significance based on ordinary two-way ANOVA. Data represented as mean ± standard deviation and *p < 0.05, **p < 0.01, ***p < 0.001, ****p < 0.0001 for all statistics.

### ΔNp63^KO^ intrapulmonary p63^+^ progenitors fail to migrate from airways

In addition to the attenuated dysplastic alveolar remodeling response in Sox2^ΔNp63KO^ mice, we noted that Sox2^ΔNp63KO^ mice had an approximately two-fold increase in the relative percentage of Sox2- trace^+^ SPC^+^ AT2s compared to Sox2^ΔNp63WT^ mice (Fig. 2h). To address the possibility that ΔNp63^KO^ intrapulmonary p63^+^ progenitors can participate in alveolar AT2 regeneration, we utilized a p63-driven CreERT2 to simultaneously lineage trace and delete ΔNp63 specifically in intrapulmonary p63^+^ progenitors. While IP tamoxifen caused no unexpected side effects in uninjured p63^CreERT2^; RFP (p63^ΔNp63WT^) mice, we noted that administration of even a single dose of IP tamoxifen to uninjured p63^CreERT2^; RFP; ΔNp63^flox/flox^ (p63^ΔNp63KO^) mice was lethal within 5-7 days, likely due to ΔNp63 deletion causing severe defects in epithelia that rely on basal cells for homeostatic maintenance (e.g. skin) (Candi et al., 2006; Laurikkala et al., 2006; Pellegrini et al., 2001; Romano et al., 2009; Romano et al., 2012; Signoretti et al., 2005; Signoretti et al., 2000; Truong et al., 2006). To circumvent this lethality, we intranasally administered three doses of 0.025mg/g 4-hydroxytamoxifen (4OHT) sonicated into PBS to p63^ΔNp63WT^ and p63^ΔNp63KO^ mice. p63^ΔNp63KO^ mice treated with this method can survive the full tamoxifen dosing, >1 week chase period, and injury time course with minimal side effects (Fig. 3a). As expected, p63-trace^+^ regions were found at day 21 post-flu within p63^ΔNp63WT^ airways and alveoli (Fig. 3b) as previously described (Xi et al., 2017; Yang et al., 2018). In stark contrast, p63-trace^+^ cells were restricted to the airways of p63^ΔNp63KO^ mice and did not extend into the alveolar space (Fig. 3c). Quantification of the number of airway- or alveolar-localized p63-trace^+^ cells per histology slide revealed a dramatic reduction in the number of p63-trace^+^ cells in both the airways and alveoli of p63^ΔNp63KO^ mice (Fig. 3d), underscoring the near-complete abolishment of intrapulmonary p63^+^ progenitor expansion without ΔNp63. Despite this clear phenotype, we did observe occasional p63- trace^+^ cells in the airways of p63^ΔNp63KO^ mice that still expressed p63 (Supplementary Fig. 2a, c) and Krt5 (Supplementary Fig. 2b, d), indicating incomplete recombination of the ΔNp63^flox^ allele. These results demonstrate that the increased percentage of Sox2-trace^+^ SPC^+^ AT2s in p63^ΔNp63KO^ mice is not due to direct conversion of p63^ΔNp63KO^ intrapulmonary p63^+^ progenitors into AT2s. Instead, these results indicate that the lack of Krt5^+^ dysplasia in ΔNp63^KO^ mice is due to loss of proliferative or migratory functions in ΔNp63^KO^ intrapulmonary p63^+^ progenitors.

**Figure 3.**
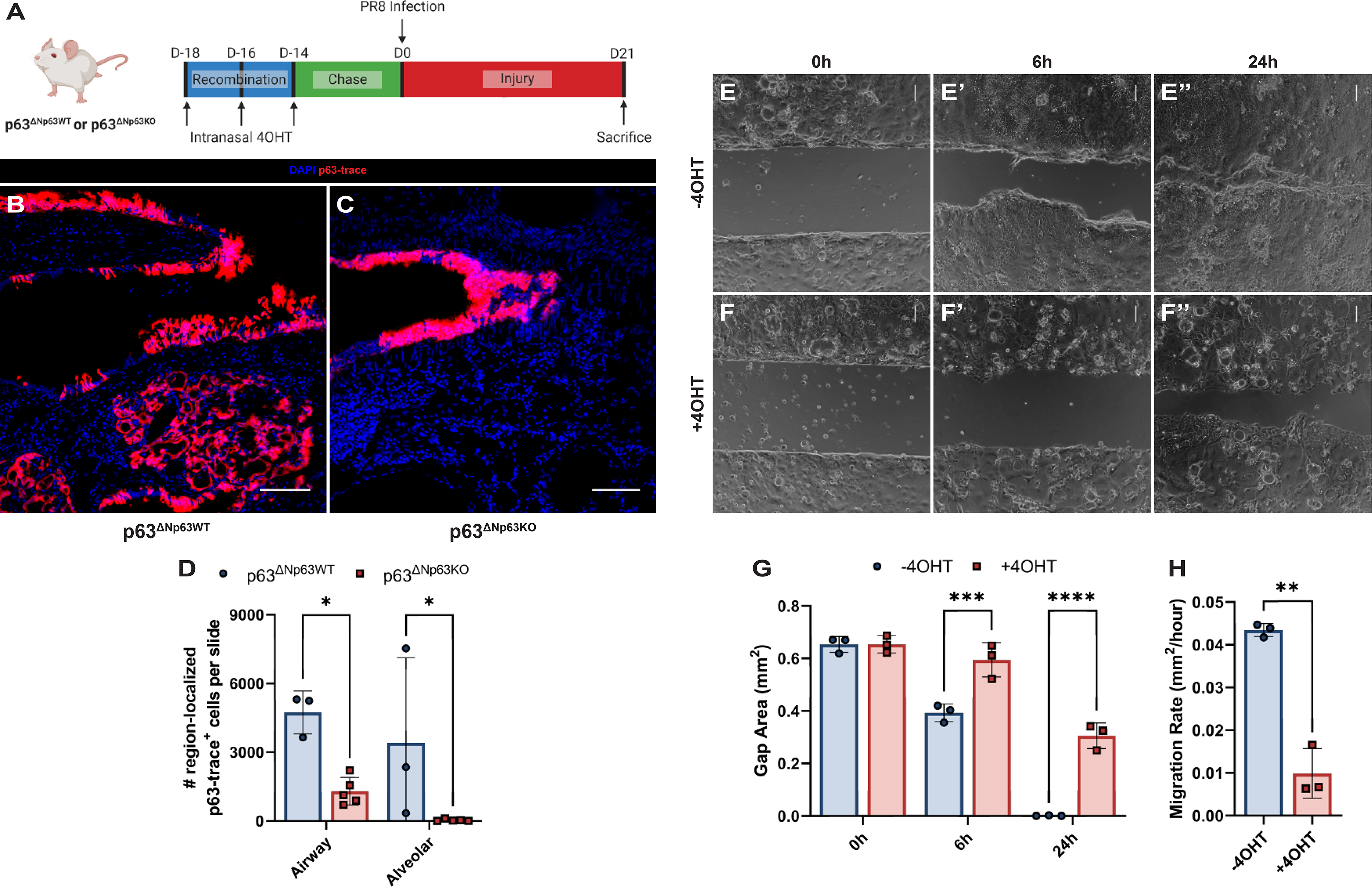
ΔNp63^KO^ intrapulmonary p63^+^ progenitors fail to migrate from airways *in vivo*. (a) Experimental timeline of p63-mediated ΔNp63 deletion. (b) p63-trace^+^ cells contribute to both airway and alveolar repair in p63^ΔNp63WT^ mice. Scale bar = 100um. (c) p63-trace^+^ cells do not contribute to alveolar repair in p63^ΔNp63KO^ mice, remaining only in the airways. Scale bar = 100um. (d) Quantification of p63-trace^+^ cells in either the airways or alveoli in p63^ΔNp63WT^ or p63^ΔNp63KO^ mice. Significance based on mixed-effects analysis. (e-f’’) Brightfield images of migration assay at (e, f) 0h, (e’, f’) 6h, and (e’’, f’’) 24h timepoints. Scale bars = 100um. (g) Quantification of gap area closure from migration experiment shows that while -4OHT-treated monolayers can close the gap by 24h post-insert removal, +4OHT- treated (ΔNp63^KO^) monolayers do not fully close the gap. Significance based on ordinary two-way ANOVA. (h) Comparison of area migration rate in mm^2^/hour, revealing that +4OHT-treated (ΔNp63^KO^) intrapulmonary p63^+^ progenitors migrate ∼four times slower than -4OHT-treated (ΔNp63^WT^) intrapulmonary p63^+^ progenitors. Significance based on Welch’s t-test. Data represented as mean ± standard deviation and *p < 0.05, **p < 0.01, ***p < 0.001, ****p < 0.0001 for all statistics.

Recognizing that the failure of ΔNp63^KO^ mice to mount a dysplastic remodeling response could be caused by attenuated migration from the airways, we assessed the impact of ΔNp63 deletion on basal-like cell migration *in vitro*. Adding the basal cell-specific cell surface marker lymphocyte antigen 6 family member D (Ly6d) (Supplementary Fig. 3 a, b) to previously established sort schemes used to isolate airway epithelial cells (Chapman et al., 2011; Vaughan et al., 2015), we were able to sort enriched intrapulmonary p63^+^ progenitors from day 14 post-flu non-tamoxifen-treated Krt5^CreERT2^; RFP; ΔNp63^flox/flox^ (Krt5^ΔNp63KO^) mice. These cells allow for inducible deletion of ΔNp63 *in vitro* via the addition/withholding of 4OHT. At 6 hours following insert removal, DMSO-treated monolayer gap areas were reduced by ∼40%, with gap areas fully closed by 24 hours (Fig. 3e-e’’, g). In contrast, 4OHT- treated monolayer migration was significantly impeded, with a gap area reduction of only ∼9% at 6 hours and without ever fully closing the gap by 24 hours (Fig. 3f-f’’, g). Comparing the derived rate of gap area closure over the initial 6-hour window, we found that 4OHT-treated monolayers migrate at only ∼1/4-1/5 of the rate of their DMSO-treated counterparts (Fig. 3h), substantiating our hypothesis that ΔNp63^KO^ intrapulmonary p63^+^ progenitors are unable to execute a normal migratory program and thus cannot participate in dysplastic alveolar remodeling following *in vivo* deletion.

### Pro-alveolar histone modifications and transcriptional changes occur with ΔNp63^KO^

p63 inhibition/deletion causes precocious differentiation of basal cells in a variety of epithelial tissues into terminally differentiated cells of that tissue (Carroll et al., 2006; Haas et al., 2019; Laurikkala et al., 2006; Romano et al., 2012; Truong et al., 2006; Yang et al., 2018). p63 directly promotes the basal cell program and actively enforces basal cell identity transcriptionally (Carroll et al., 2006; Chakrabarti et al., 2014; Ihrie et al., 2005; Romano et al., 2009; Romano et al., 2012; Signoretti et al., 2005) and epigenetically (Kouwenhoven et al., 2015; Lin-Shiao et al., 2019; Mardaryev et al., 2014; Qu et al., 2019; Santos-Pereira et al., 2019). Reasoning that the observed decrease in migratory capacity could be due to terminal differentiation into lung epithelial cells via loss of epigenetic reinforcement and decreased transcriptional regulation, we surveyed the epigenetic and transcriptional landscape of ΔNp63^KO^ intrapulmonary p63^+^ progenitors, hypothesizing that ΔNp63 deletion might promote 1.) enrichment of active and/or loss of repressive histone marks around the genomic loci of lung identity and airway/alveolar epithelial differentiation genes, and 2.) loss of active and/or gain of repressive histone marks around the genomic loci of basal cell identity genes (Fig. 4a). We performed H3K27me3 (a histone mark associated with condensed chromatin and transcriptional repression) and H3K27ac (a histone mark associated with open chromatin and transcriptional activation) chromatin immunoprecipitation followed by quantitative RT-PCR (ChIP-qPCR) on post-flu Krt5^ΔNp63WT^ and Krt5^ΔNp63KO^ cells treated for 48 hours with 4OHT. The key lung epithelial identity transcription factor Nkx2-1 exhibited dynamic changes in epigenetic modification, with multiple genomic loci at and near its translational start site becoming simultaneously depleted for H3K27me3 and enriched for H3K27ac in Krt5^ΔNp63KO^ cells (Fig. 4b), suggesting that the chromatin around this gene could be open and poised for transcriptional activation. Other cell identity-conferring transcription factors such as the airway epithelial transcription factor Sox2 remained unchanged (Fig. 4c). Other differentiation-associated genes, e.g. the ciliated cell transcription factor Foxj1, exhibited trending but non-statistically significant depletion for H3K27me3 near its translational start site and enrichment for H3K27ac at a predicted upstream promoter in Krt5^ΔNp63KO^ cells (Fig. 4d). Given that basal cells normally differentiate into airway epithelial cells, we were surprised by the lack of changes in the epigenetic status at these crucial airway differentiation and identity loci. Considering that intrapulmonary p63^+^ progenitors can rarely differentiate into AT2s following their migration into the alveoli (Xi et al., 2017) and the increased H3K27 acetylation and decreased H3K27 trimethylation around the Nkx2-1 locus – a transcription factor critical for both airway and alveolar epithelial differentiation - we turned our attention towards changes in epigenetic marks at alveolar differentiation loci. A predicted promoter region for Etv5, an essential transcription factor for AT2 identity (Zhang et al., 2017), was significantly depleted of H3K27me3 in Krt5^ΔNp63KO^ cells (Fig. 4e), suggesting its potential poising for transcriptional activation. Multiple AT2-specific loci, including regions immediately upstream and at the translational start site of SPC (Fig. 4f) and downstream of the first exon of Lamp3 (Fig. 4g), were also significantly depleted of H3K27me3. In total, these results indicate that ΔNp63 maintains the epigenetic configuration of intrapulmonary p63^+^ progenitors, and the deposition of histone marks associated with transcriptional activation and removal of histone marks associated with transcriptional repression at critical lung epithelial and alveolar transcription factors along with AT2-specific genes predict that loss of ΔNp63 may allow for rewiring of the epigenetic landscape into one amenable to alveolar differentiation.

**Figure 4.**
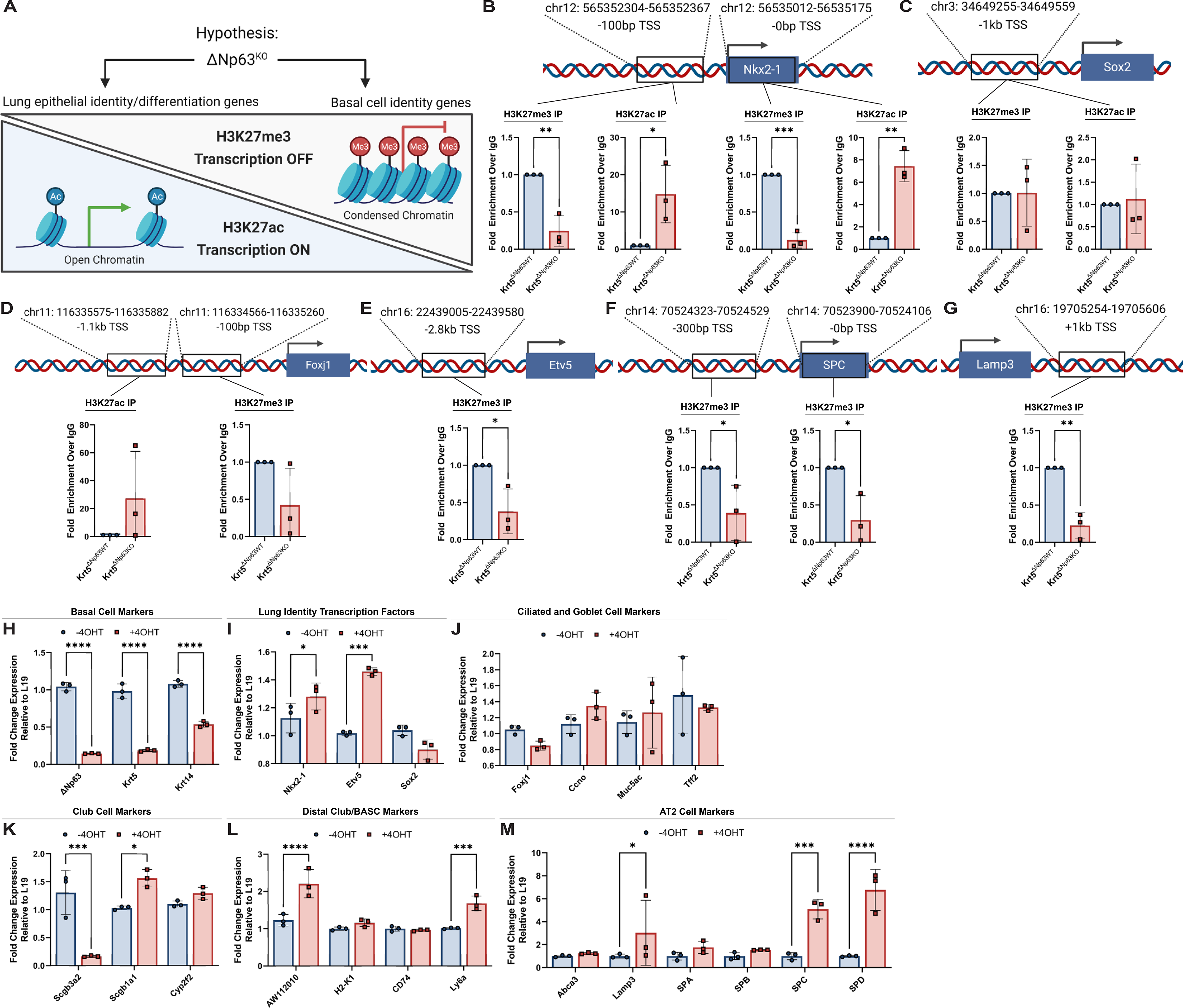
Intrapulmonary p63^+^ progenitor transcriptional and epigenetic state is regulated by ΔNp63. (a) Schematic of ChIP-qPCR experiment prediction. (b-g) ChIP for H3K27me3 or H3K27ac marks followed by qPCR (b) at and 100bp upstream of the Nkx2-1 translational start site (TSS), (c) at a predicted promoter 1kb upstream of the Sox2 TSS, (d) at a predicted promoter 1kb upstream and 100bp upstream of the Foxj1 TSS, (e) at a predicted promoter 2.8kb upstream of the Etv5 TSS, (f) at and 300bp upstream of the SPC TSS, and (g) in the first intron 1kb downstream of the Lamp3 TSS. Significance based on unpaired t-test. (h-m) qPCR of intrapulmonary p63^+^ progenitor monolayers. Alterations in histone deposition correlate with changes in transcriptional identity with ΔNp63^KO^, focusing on (h) basal cell markers, (i) lung identity transcription factors, (j) ciliated and goblet cell markers, (k) club cell markers, (l) distal club/BASC markers, and (m) AT2 markers. Significance based on ordinary two-way ANOVA. Data represented as mean ± standard deviation and *p < 0.05, **p < 0.01, ***p < 0.001, ****p < 0.0001 for all statistics.

These epigenetic changes prompted us to assess gene expression changes in Krt5^ΔNp63KO^ monolayers treated with DMSO (-4OHT) or 4OHT (+4OHT) for 48 hours. Consistent with the role of ΔNp63 in maintaining basal cell identity, basal cell marker genes Krt5 and Krt14 were significantly reduced upon 4OHT treatment (Fig. 4h). Supporting our ChIP-qPCR data, Nkx2-1 and Etv5 expression levels increased with 4OHT treatment while Sox2 expression exhibited a trending but non-statistically significant decrease in expression (Fig. 4i), indicating a shift towards a more distal airway or potentially alveolar fate. Indeed, expression of the multiciliated cell identity transcription factor Foxj1 and cilia element Ccno and the goblet cell markers Muc5ac and Tff2 were unchanged with 4OHT treatment (Fig. 4j), confirming that ΔNp63^KO^ intrapulmonary p63^+^ progenitors do not spontaneously commit to these airway lineages in monolayer conditions.

### ΔNp63^KO^ induces intrapulmonary p63^+^ progenitor plasticity

Considering the vast morphological, transcriptional, and anatomic differences between basal cells and AT2s, in addition to the rarity of intrapulmonary p63^+^ progenitor-to-AT2 differentiation *in vivo*, we hypothesized that ΔNp63^KO^ intrapulmonary p63^+^ progenitors might pass through an intermediary cell type common to both lineages. Since tracheal basal cells can differentiate into club cells during airway homeostasis and repair, and distal club cells/BASCs can differentiate into AT2s, we predicted that ΔNp63^KO^ intrapulmonary p63^+^ progenitors might assume a distal club cell/BASC-like phenotype. In line with this prediction, the proximal club cell gene Scgb3a2 was downregulated while the distal Club cell gene Scgb1a1 and distal club/BASC-specific markers AW112010 and Ly6a were induced with 4OHT treatment (Fig. 4k, l). Consistent with assumption of the distal club cell/BASC-like phenotype and progenitor properties associated with it, 4OHT-treated monolayers upregulated some, but not all, mature AT2 markers SPC, SPD, and Lamp3 (Fig. 4m), reflecting the reduction of H3K27me3 deposition at the SPC and Lamp3 loci in Krt5^ΔNp63KO^ ChIP-qPCR. These findings indicate that ΔNp63^KO^ allows p63^+^ progenitor monolayers to transition toward an intermediate club/AT2-like fate, sharing transcriptional similarities with both cellular identities while forgoing the expected trajectory of airway differentiation.

Given the transcriptional shift towards an alveolar fate with ΔNp63^KO^ as monolayers, we moved to a 3D organoid culture system since these conditions are known to support higher efficiency of AT2 generation (Jacob et al., 2017; Jacob et al., 2019) and utilized alveolar differentiation media based on our previous study with some modifications (see Methods). ΔNp63 deletion via treatment with 4OHT diminished basal cell identity as expected in organoids maintained in either airway or alveolar media (Fig. 5a, e). Nkx2-1 expression was elevated with ΔNp63^KO^ in alveolar media and not airway media, while Sox2 expression was unchanged despite ΔNp63 status or media conditions (Fig. 5b, f). ΔNp63^KO^ organoid maintenance in airway media allowed for robust expression of differentiated airway cell markers, particularly those expressed in distal club cells/BASCs, including cytochrome p450 2f2 (Cyp2f2) (Fig. 5c), AW112010, CD74 (Fig. 5l), and, interestingly, the surfactant proteins SPA and SPD (Fig. 5j), perhaps reflecting the promiscuous expression of surfactant genes seen in some distal club cells/BASCs (Kathiriya et al., 2020; Liu et al., 2019; Salwig et al., 2019). Multiciliated cell markers were also upregulated, including Foxj1 and Ccno (Fig. 5d). In contrast, airway epithelial differentiation was minimal in alveolar media conditions as multiciliated cell gene expression remained unchanged with ΔNp63^KO^ (Fig. 5h), reinforcing a model wherein differentiation of these cells is dependent both upon p63 expression and media conditions. ΔNp63^KO^ organoids in alveolar media conditions still maintained some semblance of the distal club/BASC transcriptional signature with elevated expression of Cyp2f2 and CD74 in conjunction with significantly reduced expression of proximal club marker Scgb3a2 (Fig. 5g, k). ΔNp63^KO^ organoids in alveolar media conditions appeared fully differentiated into AT2s, evidenced by heightened expression of the AT2-specific surfactant proteins SPB, SPC, and SPD as well as Lamp3 (Fig. 5l). Immunostaining reflects this conversion of ΔNp63^KO^ intrapulmonary p63^+^ progenitor organoids to AT2s when maintained in alveolar media, as Krt5 protein expression lowers and becomes disorganized with 4OHT induction and is replaced with strong, punctate SPC staining (Fig. 5m-p). Subjecting ΔNp63^KO^ organoids to airway or alveolar media conditions revealed the plasticity afforded by ΔNp63 deletion, confirming that ΔNp63^KO^ intrapulmonary p63^+^ progenitors adopt increased plasticity, similarly to distal club/BASCs, and can subsequently be coaxed towards either airway or alveolar differentiation.

**Figure 5.**
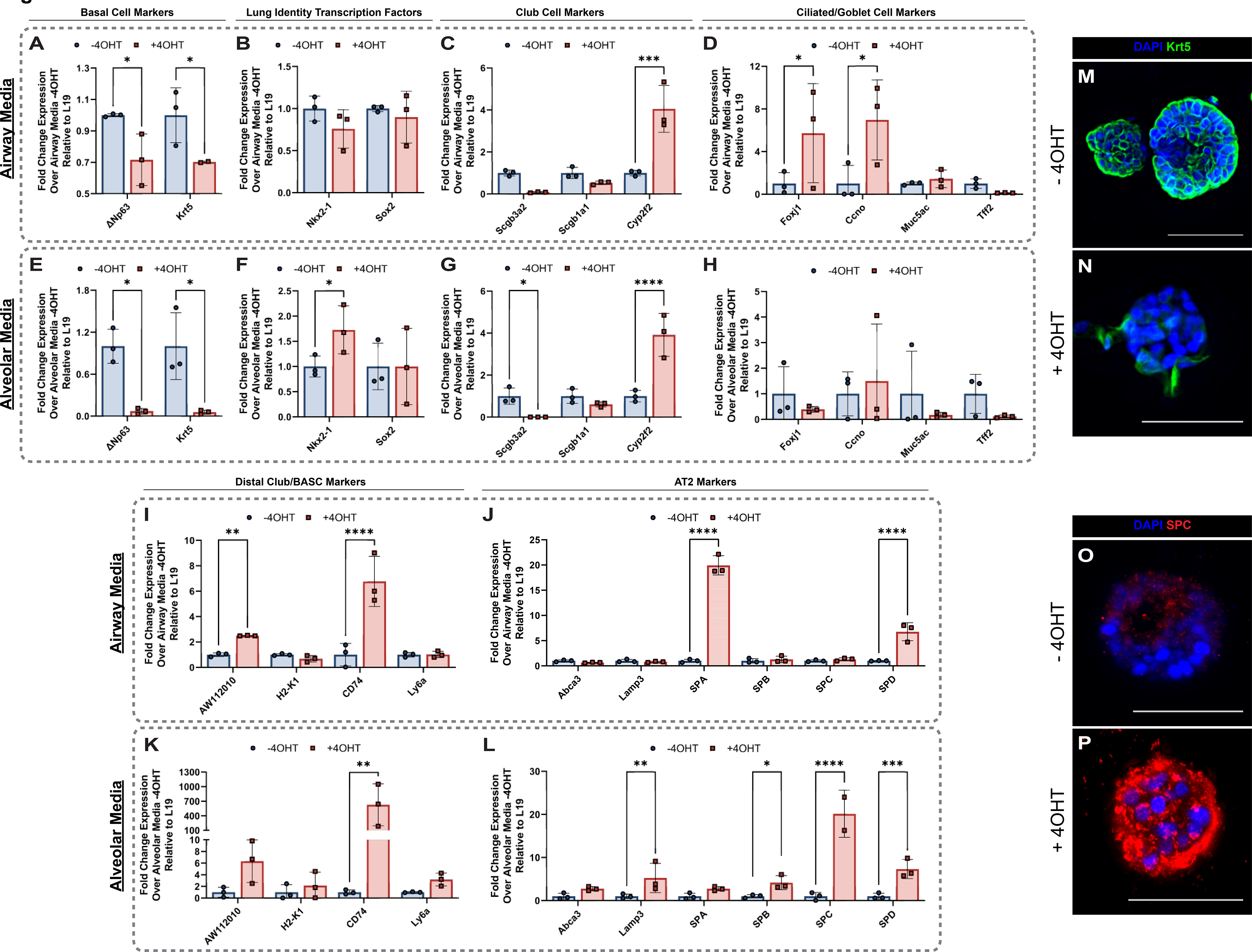
Induced plasticity in ΔNp63^KO^ intrapulmonary p63^+^ progenitors can be directed towards airway or alveolar epithelial fates. (a-l) Side-by-side comparisons of transcriptional changes in -4OHT-treated (ΔNp63^WT^) versus +4OHT-treated (ΔNp63^KO^) intrapulmonary p63^+^ progenitor organoids cultured in either (a-d, i-j) airway media or (e-h, k-l) alveolar media. (a, e) Basal cell markers are downregulated in both airway and alveolar media with ΔNp63^KO^. (b, f) Lung identity transcription factor expression remains unchanged in airway media with ΔNp63^KO^ while Nkx2-1 is upregulated in alveolar media with ΔNp63^KO^. (c, g) The club cell marker Cyp2f2 is upregulated in airway media with ΔNp63^KO^, with an enhanced shift towards a distal club-like fate evidenced by repression of Scgb3a2 in alveolar media with ΔNp63^KO^. (d, h) Ciliated cell markers are upregulated with ΔNp63^KO^ in airway media while ciliated and goblet cell markers remain unchanged with ΔNp63^KO^ in alveolar media. (i, k) Distal club/BASC markers are trending/significantly upregulated with ΔNp63^KO^ in airway and alveolar media. (j, l) The shared AT2/distal club/BASC markers SPA and SPD are upregulated in airway media with ΔNp63^KO^, while additional AT2 markers are also upregulated in alveolar media with ΔNp63^KO^. (m-p) Representative immunofluorescence images of organoids grown in alveolar media ±4OHT. Significance based on ordinary two-way ANOVA. (m, n) The basal cell marker Krt5 is downregulated and disorganized with 4OHT-treatment (ΔNp63^KO^), indicating loss of basal cell fate. (o, p) The AT2 marker SPC is upregulated with 4OHT-treatment (ΔNp63^KO^), indicating acquisition of alveolar fate. Scale bar = 50um. Data represented as mean ± standard deviation and *p < 0.05, **p < 0.01, ***p < 0.001, ****p < 0.0001 for all statistics.

Given that p63 deletion in pro-alveolar media promoted differentiation into cells provocatively similar to AT2s, we attempted to assess the effects of ΔNp63 deletion *in vivo* after dysplastic expansion. However, in spite of many attempts with different Cre drivers and routes of tamoxifen delivery, we were unable to achieve ΔNp63 deletion in the post-flu dysplastic regions with either intraperitoneal or intranasal tamoxifen delivery (Supplementary Fig. 4a-d). Attempts to increase recombination by raising the dose of tamoxifen or 4OHT invariably resulted in animal death as earlier described. Future studies utilizing novel CreER lines which allow for specific deletion in lung p63^+^ cells will be necessary to assess the fate of dysplastic cells *in vivo* upon ΔNp63 deletion.

## Discussion

The airway and alveolar repair pathways are carried out by well-defined, distinct progenitor populations: *bona fide* basal cells are responsible for proximal airway homeostasis and repair (Rock et al., 2009), whereas euplastic alveolar regeneration is carried out by AT2s and distal club/BASCs to generate more AT2s and AT1s (Barkauskas et al., 2013; Evans et al., 1973; Kathiriya et al., 2020; Kim et al., 2005; Xi et al., 2017). In severe injury, an additional pathway of dysplastic remodeling emerges, employing basal-like intrapulmonary p63^+^ progenitors to form an ectopic airway epithelium in the alveolar compartment that does not appreciably contribute to functionally beneficial regeneration (Vaughan et al., 2015; Xi et al., 2017; Yang et al., 2018). While this paradigm holds across a variety of injuries, recent studies highlight several caveats. For example, genetic ablation of basal cells in the trachea induces committed club cells to de-differentiate into basal cells, which can then respond to airway insults as endogenous basal cells would (Tata et al., 2013). Similarly, differentiated AT1s can assume progenitor functions with pneumonectomy injury, hyperoxic conditions in young mice, or Yap/Taz deletion and differentiate into AT2s (Jain et al., 2015; Penkala et al., 2021). In humans, this plasticity appears to be bidirectional, as AT2s can be driven to aberrantly transdifferentiate into basal cells when exposed to TGFβ signaling from altered CTHRC1^hi^ mesenchymal niche cells that arise in diseased lungs (Kathiriya et al., 2022). Our study provides further nuance to these models, in which injury-induced dysplastic basal-like cells can be converted into AT2s with loss of the basal cell identity transcription factor ΔNp63. While these findings challenge some paradigms re. lineage restriction and terminal differentiation, they also highlight the extreme conditions under which induced cellular plasticity occurs and provide novel mechanisms that can be leveraged to modulate pulmonary repair.

Despite the detrimental nature of long-term alveolar inhabitation by dysplastic tissue, alveolar re-epithelialization by intrapulmonary p63^+^ progenitors is thought to be critical for the short-term survival of individuals with severe injuries due to the rapid restoration of pulmonary barrier function and structural integrity that these cells provide, further evidenced by our migration studies (Fig. 3e-h). Interestingly, our data suggest that pulmonary injury is nevertheless still survivable in the absence of dysplastic repair, as mice that lose ΔNp63 in either the broad airway epithelium or specifically in intrapulmonary p63^+^ progenitors survive influenza injury, recover fully, and live until the experimental endpoint, yet exhibit no dysplastic remodeling. Additionally, our data indicate no obvious compensatory mechanism by which other airway- or non-airway-derived progenitors besides pre-existing p63^+^ progenitors can engage in dysplastic remodeling. Some studies suggest that failure to mount a dysplastic remodeling response is detrimental to the recovery of lung function, as genetic ablation of dysplastic cells in influenza-injured mice causes persistent inflammation and fibrosis as well as failure to recover normal blood oxygen saturation levels (Zuo et al., 2015). Given that dysplastic bronchiolization with basal-like cells appears to be evolutionarily conserved (Kanegai et al., 2016;

Vaughan et al., 2015), we predict that it does indeed provide some survival benefit, though proving this will require precise control of injury levels and cell type-specific ablation techniques outside the scope of what is currently achievable. Given that ΔNp63 deletion prior to injury prevents dysplastic repair from occurring while other studies deplete dysplastic cells after their injury-induced expansion, it is possible that the immune and fibrotic responses elicited in these distinct scenarios differ as well. Indeed, crosstalk between the dysplastic epithelium and the immune and mesenchymal responders to lung injury have been the focus of recent attention, and there is mounting evidence that Krt5^+^ cells can recruit pathological immune and mesenchymal subpopulations to the injured lung (Cassandras et al., 2020; Choi et al., 2020; Habermann et al., 2020; Kathiriya et al., 2022; Strunz et al., 2020; Wu et al., 2021). In light of this, absence of the dysplastic epithelium could impair recruitment of these pathologic, pro-fibrotic, and pro-inflammatory populations, which might in-and-of itself be beneficial to lung repair and allow other epithelial progenitors better able to respond to injury. This could in turn provide a potential explanation for how loss of such a substantial epithelial population in our model could ultimately benefit the injured lung, allowing for injury survival.

The mechanisms by which cells acquire, maintain, and alter their identity are critical to understanding how and why progenitor populations respond to injury and contribute to diseases such as fibrosis, chronic inflammation, and cancer (Blanco et al., 2021; Jiang et al., 2017; Kobayashi et al., 2020; Strunz et al., 2020). Our data provide a novel mechanism for ΔNp63 in maintaining the basal cell identity of intrapulmonary p63^+^ progenitors, in which ΔNp63 restricts intrapulmonary p63^+^ progenitor fate by maintaining chromatin in a repressive state at loci required for distal airway and alveolar differentiation. It is likely that ΔNp63 exerts its epigenetic control over these loci via cooperation with as-of-yet identified lung tissue- and cell-specific transcription factors, histone modifiers, and chromatin remodelers as it does in other epithelial tissues (Fessing et al., 2011; Lin-Shiao et al., 2019; Qu et al., 2018). Identification of these epigenetic factors may represent promising druggable targets for *in vivo* directed differentiation or “differentiation therapy” to either make intrapulmonary p63^+^ progenitors less stable and/or less proliferative, or to directly convert p63^+^ progenitors into functionally beneficial AT2s and AT1s.

Adding to the model in which ΔNp63 maintains basal cells in an undifferentiated stem-like state and loss of ΔNp63 leads to terminal differentiation, our data show that the differentiation trajectory of ΔNp63^KO^ airway basal cells can be modulated depending on the surrounding environment. Our observed induction of club and ciliated cell genes with ΔNp63 deletion in organoids maintained in airway media is concordant with previous studies, paralleling both the differentiation trajectory of embryonic p63^KO^ airway basal cells (Yang et al., 2018) and adult club cells when basal cells are genetically ablated (Tata et al., 2013). We highlight that 3D “organoid” conditions appear to be an important factor in basal cell differentiation, as it was only in organoid conditions - in contrast to 2D monolayers - that ΔNp63^KO^ intrapulmonary p63^+^ progenitors expressed the mature club and ciliated cell genes Cyp2f2, Foxj1, and Ccno. On the other hand, ΔNp63^KO^ intrapulmonary p63^+^ progenitors express mature AT2 genes such as Lamp3, SPB, SPC, and SPD as opposed to airway cell markers when maintained in organoid conditions in a pro-alveolar media formulation. Interestingly, ΔNp63^KO^ intrapulmonary p63^+^ progenitor organoids in either media environment maintain expression of a core set of club cell and distal club/BASC genes, including Cyp2f2, CD74, and SPD. This distal club/BASC-like state is more pronounced in ΔNp63^KO^ intrapulmonary p63^+^ progenitor organoids in airway media than alveolar media, likely due to the latter’s ability to further differentiate these cells into AT2s downstream of this distal club/BASC-like phenotype. Given the conservation of this state following ΔNp63 deletion in both environments, we propose that the club/distal club/BASC-like phenotype constitutes a “ground state” for an intrapulmonary p63^+^ progenitor that loses its basal cell identity. This model parsimoniously allows ΔNp63^KO^ intrapulmonary p63^+^ progenitors to exist in an intermediate state along the pulmonary epithelial cell hierarchy, poised towards further airway or alveolar differentiation depending on encountered signals. Whether this is a discrete, definable state *in vivo* or rather represents a relative and dynamic position on a differentiation spectrum remains to be determined.

Histone modifications at alveolar identity and differentiation loci, including the crucial AT2 development and differentiation transcription factors Nkx2-1 and Etv5 (Little et al., 2019; Little et al., 2021; Zhang et al., 2017; Zhou et al., 2008) and AT2-specific genes themselves, changes dramatically with ΔNp63 knockout. Interestingly, we found few changes in airway histone composition, which likely reflects the innate competence of intrapulmonary p63^+^ progenitors to differentiate into airway lineages, a homeostatic function of upper airway basal cells. Considering that differentiation into alveolar cells is a more dramatic shift in identity which rarely happens *in vivo*, ΔNp63^KO^ intrapulmonary p63^+^ progenitors likely require larger-scale epigenetic rewiring and chromatin remodeling to proceed down this differentiation path. This further indicates that ΔNp63 actively maintains chromatin in a configuration only amenable to airway differentiation and that the rare *in vivo* event of intrapulmonary p63^+^ progenitor differentiation into AT2s requires suppressed ΔNp63 expression levels. The intracellular mechanisms or extracellular signals that reduce ΔNp63 expression, thereby altering chromatin organization and allowing for plasticity, remain unknown and warrant further study.

A fundamental limitation of this study was the difficulty in recombining the ΔNp63^flox^ allele *in vivo* with basal cell-specific recombinases and the lethality that attempting to do so caused. While other groups report achieving ΔNp63 deletion by giving systemic tamoxifen in tandem with basal cell-specific or even ubiquitous Cre recombinases (Kumar et al., 2020; Min et al., 2020), we were only to able to achieve appreciable ΔNp63 deletion while circumventing lethality by using less broadly-expressed Cre recombinases or by delivering 4OHT directly to the lung before injury. In our hands, neither systemic tamoxifen nor intranasal 4OHT delivery combined with the basal cell-specific Krt5^CreERT2^ to delete ΔNp63 after injury were sufficient to fully recombine the alleles (Supplementary Fig. 4). While failure to detect appreciable ΔNp63 deletion could be due to the mice succumbing before full recombination of both ΔNp63 alleles could occur, this could also point to biologically relevant aspects of fully formed dysplastic pods, namely that 1.) They are likely poorly vascularized, which could explain their association with hypoxia (Xi et al., 2017) and why systemic tamoxifen was unable to reach the Krt5^+^ pods in a large enough quantity or for a sustained enough period of time to recombine the floxed alleles, and 2.) They might not be directly connected to the rest of the intrapulmonary airways (i.e. they exist partially as cysts), which is also supportive of why Krt5^+^ pods have adapted to thrive under hypoxic conditions and intranasal 4OHT was unable to induce full recombination. While our data strongly support the conclusion that ΔNp63 deletion induces intrapulmonary p63^+^ progenitor plasticity and that further culturing of ΔNp63^KO^ intrapulmonary p63^+^ progenitors in alveolar organoid conditions further promotes differentiation into AT2s *in vitro*, further studies using intrapulmonary p63^+^ progenitor-specific recombinases or alternative deletion methods such as CRISPR/Cas9 recombineering will be necessary to validate this finding *in vivo*.

This study identifies ΔNp63 as a critical driver of dysplastic alveolar remodeling and elucidates novel mechanisms that promote intrapulmonary p63^+^ progenitor plasticity, allowing for their differentiation into AT2s. The dramatic regulation of dysplastic remodeling and epigenetic control of lung identity by ΔNp63 expands on our knowledge of lung progenitor cell biology, plasticity, and roles in injury repair, providing nuance to the interconnected lung epithelial cell hierarchy and the ability of vastly different lung progenitors to converge upon similar cell states. Additionally, we demonstrate that heightened ΔNp63 expression is associated with diseased states in both mice and humans. The novel mechanisms of lung repair and cell identity governance identified in this work also highlight exciting avenues for the development of therapies to treat human patients afflicted with a wide array of severe pulmonary injuries and diseases that involve dysplastic alveolar remodeling, including ILD, ARDS/DAD, and COVID-19.

## Methods

### Animals and treatment

8- to 10-week-old mice were used for all experiments with males and females in roughly equal proportions. Experimenters were not blinded to mouse age or sexes. Ai14tdTomato (Gt(ROSA)26Sor^tm14(CAG-tdTomato)Hze^) (Madisen et al., 2010), ΔNp63^flox^ (Chakravarti et al., 2014; Su et al., 2009), Sox2^CreERT2^ (*Sox2^tm1(cre/ERT2)Hoch^*) (Arnold et al., 2011), Krt5^CreERT2^ (Van Keymeulen et al., 2011), and p63^CreERT2^ (Lee et al., 2014) mice have been previously described. All studies were approved by the University of Pennsylvania’s Institutional Animal Care and Use Committees, protocol 806262, and followed all NIH Office of Laboratory Animal Welfare regulations. Genotyping primers for all transgenic mice used are in Table 1.

**Table 1.**
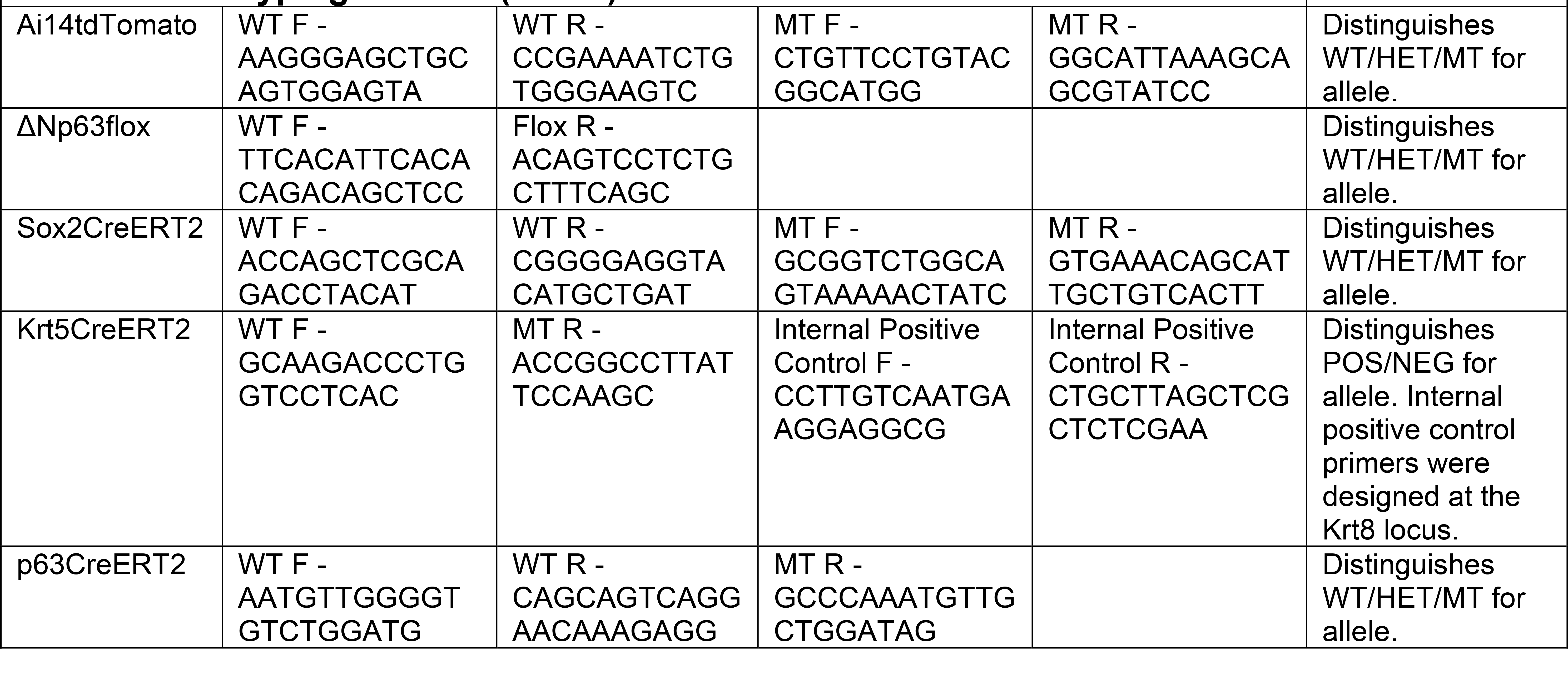
Genotyping Primers (5’-3’)

### Influenza virus propagation and infection

Influenza A/H1N1/Puerto Rico/8/34 (PR8) was propagated in embryonated chicken eggs while IAV-Cre was propagated in MDCK cells, both according to established protocols (Xue et al., 2016). Virus titration by TCID50 for infectivity titer was performed in MDCK cells for both viral strains (Xue et al., 2016). For infection with either PR8 or IAV-Cre, mice were first anesthetized using 3.5% isoflurane in 100% O2 via an anesthesia vaporizer system. Mice were intranasally administered (2 * bodyweight in grams) – 5 U TCID50 (ex. A 25g mouse gets (2 * 25g) - 5U = 45U) PR8 by pipetting 30uL of virus dissolved in phosphate-buffered saline (PBS) onto the nostrils of anesthetized mice in visually confirmed agonal breathing. Since IAV-Cre takes a higher dosing to achieve comparable injury (Hamilton et al., 2016; Heaton et al., 2014), anesthetized mice in agonal breathing were administered (8 * bodyweight in grams) TCID50 U (ex. A 25g mouse gets (8 * 25g) = 200U) IAV-Cre intranasally. Only infected mice that lost ≥10% of their starting body weight by day 9 post-flu were considered to be adequately infected with either strain and were used for all experiments involving influenza infection. Mouse weights were tracked throughout infection until the experimental endpoint. All mice that received PR8 for histology experiments were sacrificed at day 21 post-flu. All mice that received IAV-Cre for histology experiments were sacrificed at day 28 post-flu. All mice that received PR8 for cell sorting were sacrificed at day 14 post-flu unless otherwise indicated.

### Intranasal and intraperitoneal tamoxifen administration

For intraperitoneal tamoxifen experiments, mice were injected with a single 0.25mg/g dose of intraperitoneal (IP) tamoxifen dissolved in 50uL corn oil into the intraperitoneal cavity and allowed ≥1 week for allele recombination and tamoxifen clearance prior to influenza infection. For intranasal 4OHT experiments, 0.025mg/g (E/Z)-4-hydroxytamoxifen (17308, Cayman Chemical) of a 100mg/mL stock in DMSO was sonicated into 30uL PBS at 4°C using an S220 Focused-Ultrasonicator (Covaris) with sonication at peak power = 140, duty factor = 5, 200 cycles/burst for 60 seconds followed by 30 second delay for 20 cycles. Mice were then anaesthetized and intranasally administered the sonicated solution according to the procedure in “Influenza infection”. Mice treated with intranasal 4OHT were given three doses every other day over the course of six days and allowed a two-week chase for allele recombination and tamoxifen clearance prior to influenza infection.

### Pulse oximetry

Repeated measurements of peripheral oxygen saturation (SpO2) were taken using a MouseOx Plus Rat & Mouse Pulse Oximeter and a MouseOx small collar sensor (Starr Life Sciences Corp.). Mice were shaved around the neck and shoulders where the collar sensor sits at least one day prior to the initial reading. Recordings were taken using MouseOx Premium Software (Starr Life Sciences Corp., Oakmont, PA, USA). Measurements were taken continuously for > 3 minutes at a measurement rate of 15 Hz. Measurements were imported into Microsoft Excel and all readings with a non-zero Error Code were discarded. The average of these error-free readings was used to calculate the SpO2 reading for each mouse for each given time point.

### Lung tissue preparation for immunostaining

Following euthanasia, lungs were inflated with 1mL of 3.2% paraformaldehyde (PFA) and incubated in 3.2% PFA for 1 hour at room temperature. Fixed lungs were then washed in multiple PBS washes over the course of 1 hour at room temperature, followed by an overnight incubation in 30% sucrose in PBS shaking at 4°C, and then a 2-hour incubation in 15% sucrose in PBS + 50% OCT compound (Fisher HealthCare) at room temperature. Finally, fixed lungs were randomly oriented and embedded in OCT by flash freezing with dry ice and ethanol.

### Immunostaining

All protocols were used identically for both mouse and human tissue. 7um sections were cut on a Leica CM3050 S Research Cryostat (Leica Biosystems). Tissue sections were further fixed for 5 minutes in 3.2% PFA, rinsed three times with PBS, and blocked in blocking solution (PBS + 1% bovine serum albumin (Affymetrix) + 5% normal donkey serum (Jackson Immuno Research) + 0.1% Triton X-100 (Millipore Sigma) + 0.02% sodium azide (Millipore Sigma)) for > 30 minutes. Slides were incubated in primary antibodies (listed below) in blocking solution overnight at 4 °C. Slides were then washed three times with PBS + 0.1% Tween-20 (Millipore Sigma) and subsequently incubated with secondary antibodies (listed below) for > 2 h at room temperature. Slides were then washed once more with PBS + 0.1% Tween-20 prior to incubation in 1uM DAPI (Life Technologies) for 5 minutes, rinsed with PBS, and mounted with either Prolong Gold (Life Sciences) or Fluoroshield (Millipore Sigma). The following primary antibodies were used: rabbit anti-SPC (1:2000, Millipore), rat anti-RAGE (1:500, R&D, clone 175410), rabbit anti-Krt5 (1:1000, BioLegend, clone Poly19055), chicken anti-Krt5 (1:500, BioLegend, clone Poly9059), goat anti-Scgb3a2 (1:200, R&D, clone AF3465), rabbit anti-p63α (1:500, Cell Signaling, clone D2K8X), rabbit anti-ΔNp63 (1:100, BioLegend, clone Poly6190), and rat anti-Ki67 (1:1000, Invitrogen, clone SolA15). The following secondary antibodies were used: Alexa Fluor™ 488- conjugated donkey anti-rabbit (1:1000, Thermo Fisher Scientific), Alexa Fluor™ 488 donkey anti-rat (1:1000, Thermo Fisher Scientific), Alexa Fluor™ 488-conjugated donkey anti-goat (1:1000, Thermo Fisher Scientific), donkey anti-chicken Alexa Fluor™ 488 (1:500, Jackson Immuno Research), Alexa Fluor™ 647-conjugated donkey anti-rabbit (1:1000, Thermo Fisher Scientific), Alexa Fluor™ 647- conjugated chicken anti-rat (1:1000, Thermo Fisher Scientific), Alexa Fluor™ 647-conjugated donkey anti-goat (1:1000, Thermo Fisher Scientific), and donkey anti-chicken Alexa Fluor™ 488 (1:500, Jackson Immuno Research). The following conjugated antibodies were used: FITC-conjugated anti-mouse Ly6d (1:100, BioLegend, clone 49-H4). All protocols and primary, secondary, and conjugated antibodies were used for both mouse and human tissue.

### Imaging

All images were taken on Leica DMi8 Microscope using a Leica DFC9000 sCMOS camera and Leica Application Suite X (LASX) software. Immunofluorescent and brightfield images were taken at either 10x, 20x, or 63x using the Z-stack and extended depth-of-field functions. Images were minimally processed prior to quantification or publication by adjusting the lookup table to lower the signal-to-noise ratio.

### Immunofluorescent lineage tracing quantification

For quantifying Sox2-traced cells in Sox2^ΔNp63WT^ and Sox2^ΔNp63KO^ mice, sections were cut and stained at three discrete levels of the tissue block separated by at least 300um each. In order to survey a large region of the lung and quantify as many airway/alveolar p63-trace^+^ expansion events as possible in p63^ΔNp63WT^ and p63^ΔNp63KO^ mice, we sampled twice the number of sections from each of three 300um levels. Only sections containing either airway or alveolar p63-trace^+^ cells were stained and quantified. If no p63-trace^+^ cells in either location were found in a given level, a new level was cut and surveyed. Cell counting was performed in LASX software and copied to Microsoft Excel for further analysis.

### Fluorescence-activated cell sorting

Lung cells were isolated by inflating lungs with 15 U/mL dispase II in HBSS (Thermo Fisher Scientific), tying off the trachea, and cutting lobes away from the mainstem bronchi. Lobes were then incubated in 15 U/mL dispase II for 45 minutes while shaking at room temperature and then mechanically dissociated by pipetting in sort buffer (SB; Dulbecco’s modified Eagle’s medium (DMEM) (Thermo Fisher Scientific) + 2% cosmic calf serum (CC; Thermo Fisher Scientific) + 1% P/S). After pelleting at 550 × g for 5 minutes at 4°C, whole-lung suspension was treated with Red Blood Cell Lysis Buffer (Millipore Sigma) for 5 minutes, pelleted, and resuspended in SB + 1:1000 DNase I (Millipore Sigma) for a 45-minute recovery period shaking at 37°C. Whole-lung suspension was then repelleted and resuspended in SB + 1:50 TruStain FcX™ antibody (anti-mouse CD16/32, BioLegend) for a 10-minute blocking period at 37°C. To sort post-flu intrapulmonary p63^+^ progenitors from Krt5^CreERT2^ mice, non-tamoxifen-treated Krt5^ΔNp63WT^ and Krt5^ΔNp63KO^ mice that lost ≥10% bodyweight were euthanized via isoflurane overdose at day 14 post-flu, had their lungs processed as above, and were stained with APC/Cy7-conjugated rat anti-mouse CD45 antibody (1:200, BioLegend, clone 30-F11), Alexa Fluor^®^ 488-conjugated or PE/Cy7-conjugated rat anti-mouse CD326 (EpCAM) antibody (1:200, BioLegend, clone G8.8), Alexa Fluor^®^ 647-conjugated rat anti-mouse CD104 (integrin β4) antibody (1:100, BioLegend, clone 346-11A), and FITC-conjugated rat anti-mouse Ly6d (1:100, BioLegend, clone 49-H4) for 45 minutes at 4 °C, followed by a final spin-down at 550 × g for 5 minutes at 4°C. Stained cells and fluorescence minus one controls were then resuspended in SB + 1:1000 DNase + 1:1000 Draq7 (Beckman Coulter) as a live/dead stain. All FACS sorting was done on either a BD FACSJazz (BD Biosciences) or BD FACSAria (BD Biosciences) and cells were collected in round-bottom polystyrene tubes (Thermo Fisher Scientific) in 300uL DMEM + 20% CC + 2% P/S.

### Primary intrapulmonary p63^+^ progenitor culture and passaging

Post-flu intrapulmonary p63^+^ progenitors were isolated from non-tamoxifen-treated Krt5^ΔNp63WT^ and Krt5^ΔNp63KO^ mice by sorting live CD45^-^ EpCAM^+^ β4^+^ Ly6d^+^ cells from injured lungs at day 14 post-flu. Sorted intrapulmonary p63^+^ progenitors were seeded onto 70uL of Matrigel (BD) in a 96-well clear polystyrene flat-bottom microplate (Millipore Sigma) to generate organoids. Media used for airway culture conditions was based on a modified dual SMAD inhibition media utilized for the long-term culture of basal epithelial cells (Mou et al., 2016) and allowed for intrapulmonary p63^+^ progenitor growth as either organoids or monolayers, consisting of PneumaCult^™^-Ex Plus Medium (Stemcell Technologies) + 1x PneumaCult^™^-Ex Plus Supplement (Stemcell Technologies) + 1:1000 hydrocortisone stock solution (Stemcell Technologies) + 1uM A8301 (TGFβ inhibitor, Millipore Sigma) + 10uM Y-27632 (ROCK inhibitor, Cayman Chemical) + 1:5000 Primocin^®^ (Invivogen). Media used for alveolar culture conditions was based on conditions for primary mouse AT2 (C12) (Weiner et al., 2019) and human iPSC-derived AT2 (CK+DCI) (Jacob et al., 2017; Jacob et al., 2019) culture. Briefly, CK+DCI was made according to (Jacob et al., 2019) and supplemented with 50ng/mL EGF (Peprotech) + 200ng/mL Rspo1 (Peprotech) + 100ng/mL Noggin (Peprotech) + 100ng/mL FGF10 (Peprotech) + 1uM A8301 (TGFβ inhibitor) (Millipore Sigma) + 10uM Y-27632 (ROCK inhibitor) (Cayman Chemical). To passage organoids, organoids were treated with 15 U/mL dispase II in Hank’s balanced salt solution (HBSS) for 30 minutes, rinsed with 2 mM EDTA (Thermo Fisher Scientific), incubated in 0.05% trypsin (Gibco) + 2 mM EDTA in PBS for 5 min at 37°C, mechanically dissociated by pipetting 50–100 times with a p200 pipette, quenched with DMEM + 10% CC + 1% P/S, and pelleted at 550 × g for 5 minutes at 4°C. Organoid seeding density for all experiments was 10k dissociated cells/96-well plate well. To passage monolayers, monolayers were treated with 0.05% trypsin (Gibco) + 2 mM EDTA in PBS for 10 min at 37°C, quenched with DMEM + 10% CC + 1% P/S, and pelleted at 550 × g for 5 minutes at 4°C.

### Migration assay and quantification

Migration assays were performed using 2-well silicone inserts (Ibidi). Upon securing a silicone insert on a culture plate, the surface within the insert was coated with 0.5% gelatin from bovine skin (Millipore Sigma) + 5% Matrigel (BD) for 15 minutes. Non-tamoxifen-treated intrapulmonary p63^+^ progenitors from Krt5^ΔNp63KO^ mice were previously adapted to monolayer conditions by passaging organoids and plating dissociated cells on 0.5% gelatin from bovine skin + 5% Matrigel for at least one passage as monolayers prior to seeding for the migration assay. 30k intrapulmonary p63^+^ progenitors resuspended in airway media + 1uM 4OHT (Cayman Chemical) or an equivalent volume of DMSO were seeded on both sides of the silicone insert and allowed to form monolayers for 48 hours. After 48 hours, the insert was removed and images were taken immediately for the 0h timepoint. Images were also taken at 6h and 24h following insert removal. 10x magnification brightfield images were taken at each timepoint. Three discreet regions of the gap were imaged at each timepoint for each replicate of each condition (- 4OHT and +4OHT). All experiments were performed in technical triplicate with cells derived from the same original donor mouse between passages 8 and 10. Images in TIFF format were exported from LASX to FIJI. After setting the image scale, the “Polygon selection” tool was used to trace the gap and measured using the “Analyze -> Set Measurements -> Area” function in FIJI for all regions imaged for each replicate for each condition. We derived the migration rate by subtracting the area at 6h after insert removal from the area at 0h after insert removal from that same replicate and dividing by the time elapsed (6 hours) for each replicate for each condition (ex. (0h area – 6h area)/6).

### Monolayer and organoid culture qPCR experiments

For monolayer qPCR experiments, post-flu intrapulmonary p63^+^ progenitors from a non-tamoxifen-treated Krt5^ΔNp63KO^ mouse were expanded as organoids in airway media and adapted to monolayer conditions as in “Migration assay and quantification”. Monolayers were maintained in airway media for these experiments. Technical triplicates for the -4OHT and +4OHT groups were expanded in parallel. 1uM 4OHT was added to the media for all triplicates when they reached ∼80% confluence and was maintained in the media for 48 hours. Monolayer cells were harvested for RNA using the monolayer passaging method in “Primary intrapulmonary p63^+^ progenitor culture and passaging”, pelleted at 550 × g for 5 minutes at 4°C, and stored directly at -20°C until RNA extraction. For organoid qPCR experiments, post-flu intrapulmonary p63^+^ progenitors from a non-tamoxifen-treated Krt5^ΔNp63KO^ mouse were first expanded as organoids in airway media seeded on top of 70uL Matrigel. Organoids were then passaged using the organoid passaging method in “Primary intrapulmonary p63^+^ progenitor culture and passaging” and seeded at 10k cell/well in either airway media + 1uM 4OHT or an equivalent volume of DMSO or alveolar media + 1uM 4OHT or an equivalent volume of DMSO, all on top of 70uL Matrigel. Organoids were maintained in these conditions for seven days and were harvested for RNA using the organoid passaging method in “Primary intrapulmonary p63^+^ progenitor culture and passaging”, pelleted at 550 × g for 5 minutes at 4°C, and stored directly at -20°C until RNA extraction.

### Organoid histology

For organoid histology, dissociated monolayer cells were seeded inside a 50uL Matrigel droplet on a 6-well clear polystyrene flat-bottom plate (Millipore Sigma) at a density of 250 cell/uL Matrigel in the indicated media conditions + 1uM 4OHT or an equivalent volume of DMSO at the time of seeding. After culture for one week in these conditions, droplets were incubated in 1x TBS for 10 minutes, fixed with IHC Zinc Fixative (BD Biosciences) for 30 minutes, incubated in 30% sucrose in TBS shaking overnight at 4°C, and incubated for two hours in 15% sucrose in TBS + 50% OCT compound at room temperature. Finally, the fixed organoid-containing Matrigel droplets were scraped from plate with a weighing spatula and randomly oriented and embedded in OCT by flash freezing with dry ice and ethanol. Sectioning and immunostaining were performed as described in “Immunostaining”.

### Chromatin immunoprecipitation (ChIP) and ChIP-qPCR

Post-flu intrapulmonary p63^+^ progenitors between passages 8 and 10 from non-tamoxifen-treated Krt5^ΔNp63WT^ and Krt5^ΔNp63KO^ mice were expanded as monolayers on 10cm plates coated with 0.5% gelatin from bovine skin + 5% Matrigel in airway media. Technical triplicates from the donor Krt5^ΔNp63WT^ mouse and donor Krt5^ΔNp63KO^ mouse were expanded in parallel. 1uM 4OHT was added to the media for all triplicates when they reached ∼80% confluence and was maintained in the media for 48 hours. ChIP was performed using the Cell Signaling SimpleChIP^®^ Plus Enzymatic Chromatin IP Kit (Magnetic Beads) (Cell Signaling) and protocol. Briefly, monolayers were crosslinked with 1% paraformaldehyde for 10 minutes followed by nuclei isolation, chromatin sonication into 100-500bp fragments, protein/chromatin complex pulldown, and crosslink reversal. 1ug of rabbit anti-H3K27me3 (Cell Signaling, ref. 9733), rabbit anti-H3K27ac (Active Motif, ref. 39133), and rabbit anti-IgG (Cell Signaling, ref. 2729) antibody and 5ug of chromatin were used for each immunoprecipitation. Precipitated DNA was quantified by qPCR with the primer sets listed in Table 2. ChIP-qPCR primers were designed using the UCSC Genome Browser ENCODE Regulation Tracks, specifically by searching for predicted promoters/enhancers/gene bodies at selected target genes with enriched or depleted H3K27me3/H3K27ac peaks at various developmental and postnatal timepoints in the lung.

**Table 2.**
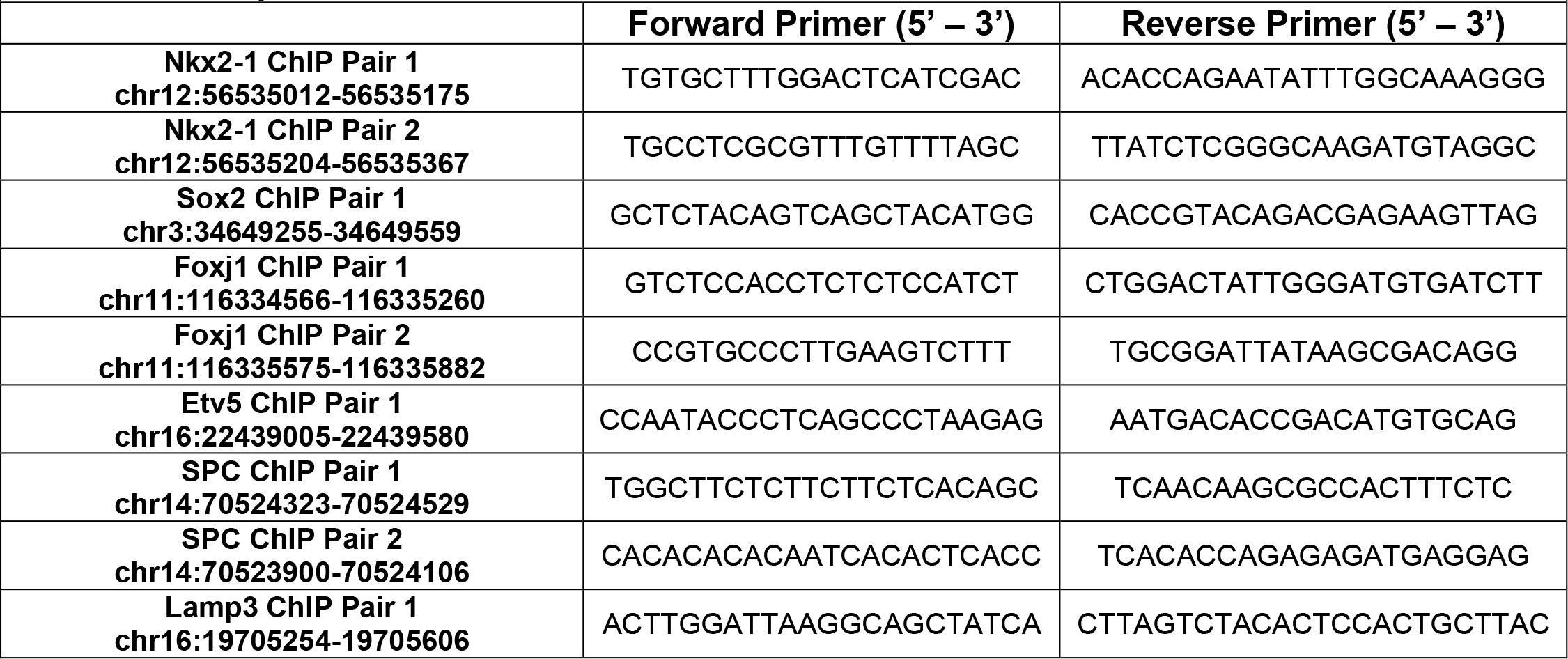
ChIP qPCR Primers

### RNA isolation, amplification, and qPCR

RNA was isolated from both monolayers and organoids using the ReliaPrep™ RNA Cell Miniprep kit (Promega). For monolayer qPCR, the amount of RNA input for cDNA synthesis was standardized within each experiment to the RNA isolate with the lowest concentration as measured by Nanodrop (Thermo Fisher Scientific). cDNA was synthesized using either iScript™ Reverse Transcription Supermix (BioRad) or High-Capacity cDNA Reverse Transcription Kit (Applied Biosystems) for monolayer experiments. For the organoid experiments, which had lower RNA yields, RNA from each sample was amplified using the Takara SMART-Seq HT Kit (Takara) according to manufacturer’s instructions and run for 20 cycles. For organoid experiments, a 1:20 dilution of amplified cDNA in deionized water was used as input in each qPCR reaction well. For injury time course qPCR of ΔNp63 expression, ≥300 live CD45^-^ EpCAM^+^ β4^+^ p63-trace^+^ cells were sorted at each indicated timepoint directly into Takara SMART-Seq HT Kit 1x reaction buffer on ice and amplified for 20 cycles. A 1:20 dilution of amplified cDNA in deionized water was used as input in each qPCR reaction well. For monolayer experiments, 2-10ng cDNA was used as input in qPCR reaction well. Expression of each gene is relative to expression of *Rpl19 (“*L19”) within that sample and expressed as fold change over the average expression relative to L19 among replicates in the -4OHT sample. qPCR was run on an Applied Biosystems QuantStudio 6 Real-Time PCR System (Thermo Fisher Scientific) with PowerUp SYBR Green Master Mix (Applied Biosystems). All primer sets are as listed in Table 3.

**Table 3.**
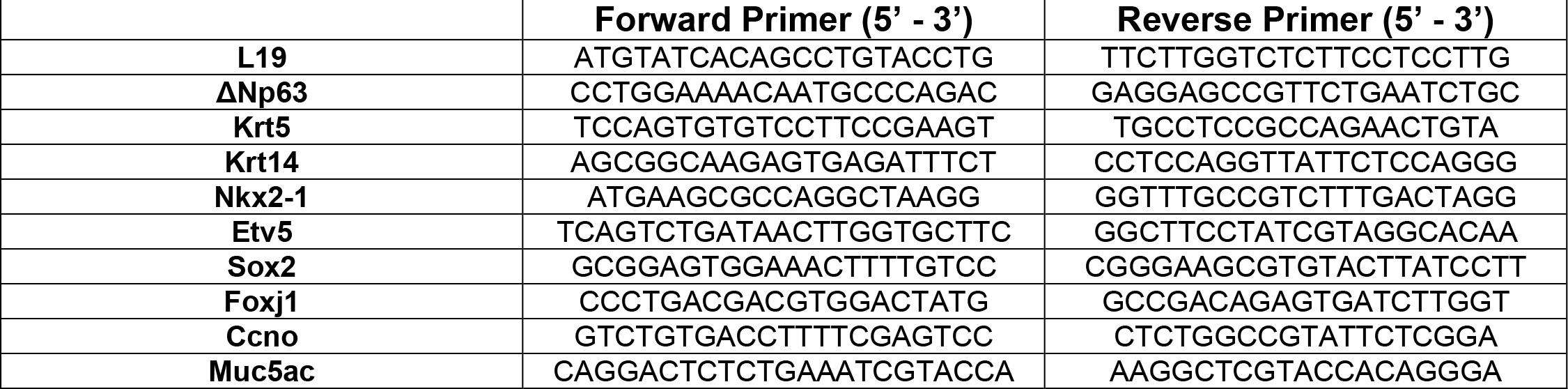
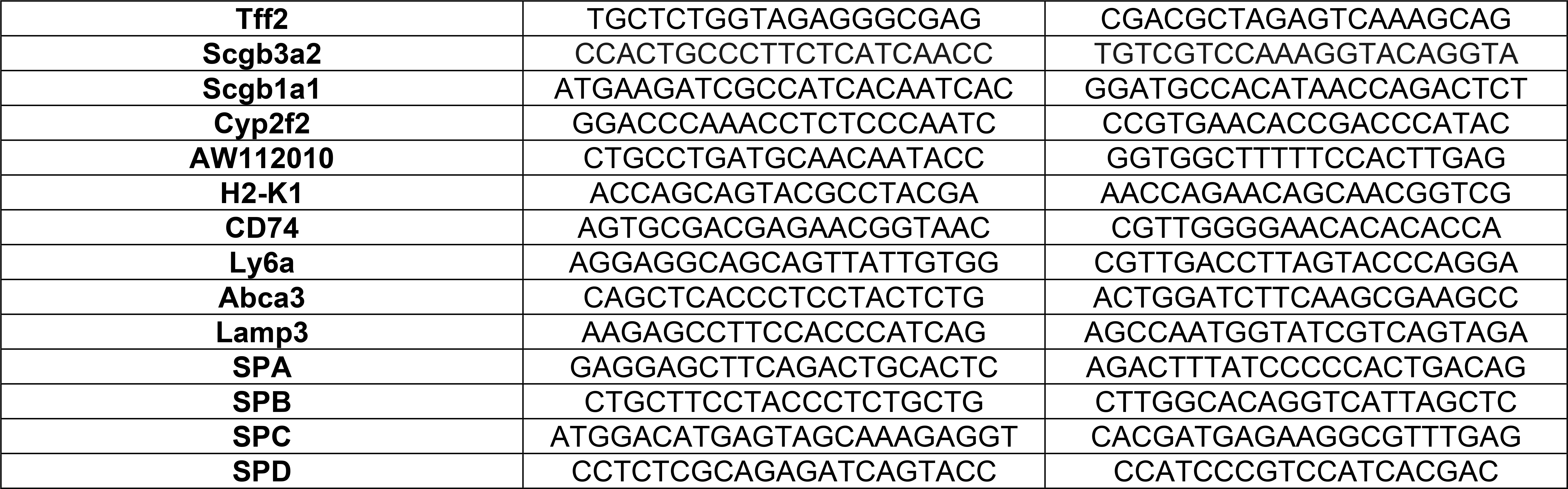
qPCR Primers

### Human tissue

Human tissue samples were obtained from either the UCSF Interstitial Lung Disease Blood and Tissue Repository (ILD samples) or the Penn Lung Biology Institute Human Lung Tissue Bank (normal lung and COVID-19 samples) and are de-identified, otherwise discarded human tissues. COVID-19 samples were from patients who previously tested positive for COVID-19 by PCR test but tested negative via PCR test prior to tissue acquisition. All COVID-19 samples were obtained from ventilated ARDS patients who underwent lung transplant, at which time tissue samples were acquired.

### Human single cell sequencing analysis

The annotated full-size RDS file from the single cell sequencing dataset generated by (Habermann et al., 2020) from healthy and IPF patients was downloaded from the Gene Expression Omnibus (GEO) under the accession code GSE135893. In R Studio using the Seurat library, cells from the clusters annotated “Basal” and “KRT5-/KRT17+” were subsetted out with the patient’s health status and TP63 expression and saved in CSV format. In Microsoft Excel, TP63 expression of the cells from each patient extracted in this way was averaged and stratified based on health status. No additional analyses, re-clustering, or *ad hoc* processing was performed on the downloaded dataset to perform this analysis.

### Statistics

Data was collected in Microsoft Excel and imported into GraphPad Prism 7 to generate all graphs and perform all statistical testing. Details of statistical test used, sample size, and definition of significance can be found in corresponding figure legends.

## Acknowledgements

We thank the Penn Lung Biology Institute Human Lung Tissue Bank, UCSF Interstitial Lung Disease Blood and Tissue Repository, and CHOP Flow Cytometry Core for their assistance in performing these studies. We also thank Dr. Montserrat Anguera and Isabel Sierra for their invaluable insight into designing the ChIP-qPCR experiment, recommendations on epigenetics bioinformatics tools, and generosity with reagents. We thank Dr. Noam Cohen for facilitating additional access to flow sorters. We thank members of the Morrisey lab for intellectual discussion. Finally, we thank BioRender for providing a platform to create the cartoons and schematics used in figures throughout this report.

---

**Supplementary Figure 1.**
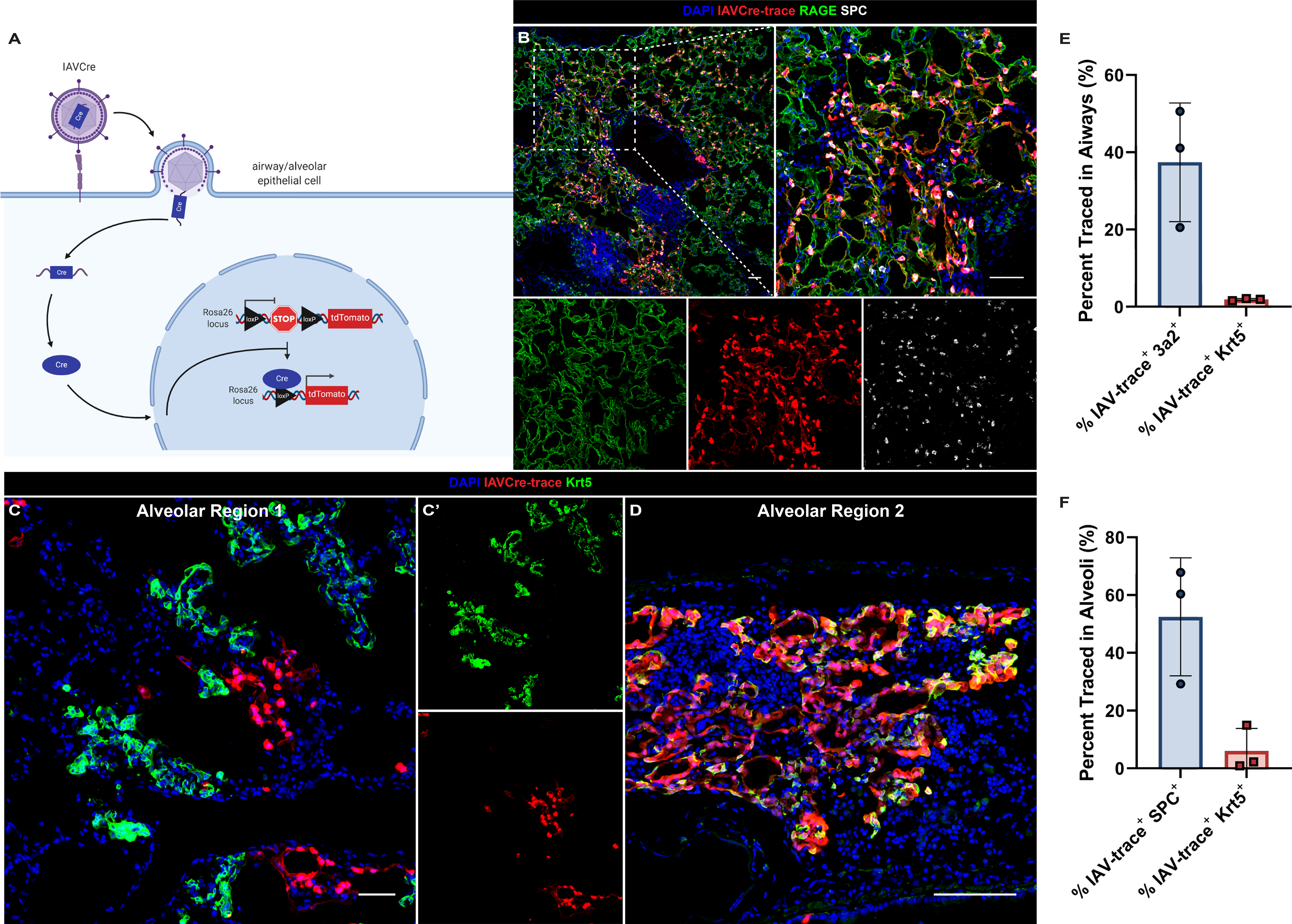
Influenza A primarily targets AT2 and club cells and rarely directly infects intrapulmonary p63^+^ progenitors. (a) Mechanism of IAV-Cre-mediated lineage tracing. IAV- Cre encodes Cre recombinase, which is taken up by actively infected cells. If infected cells harbor a lineage tracing allele such as ai14-tdTomato, the encoded Cre will excise the STOP codon, allowing for permanent tracing of the infected cell and any daughter cells that may arise from it. (b) Representative immunostaining of an IAV-Cre-traced alveolar region, which consists primarily of lineage-traced SPC^+^ AT2s and RAGE^+^ AT1s. Inset (right) is of outlined region. Separated channels of inset are below. Scale bar = 50um. (c, d) The majority of Krt5^+^ dysplastic cells are not traced by IAV- Cre, indicating that they are not directly infected by influenza. Most alveolar dysplasia (c, c’) does not overlap with lineage trace, though rare (d) regions of IAV-Cre-trace^+^ Krt5^+^ alveolar expansion can be observed (seen only in one mouse of three surveyed). (c’) is separated channels of (c). Scale bar for (c) = 50um. Scale bar for (d) = 100um. (e, f) Quantification of IAV-trace contribution to (e) Scgb3a2^+^ club cells and Krt5^+^ intrapulmonary p63^+^ progenitors in the airways and (f) SPC^+^ AT2s and Krt5^+^ dysplastic cells in the alveoli. Data represented as mean ± standard deviation.

**Supplementary Figure 2.**
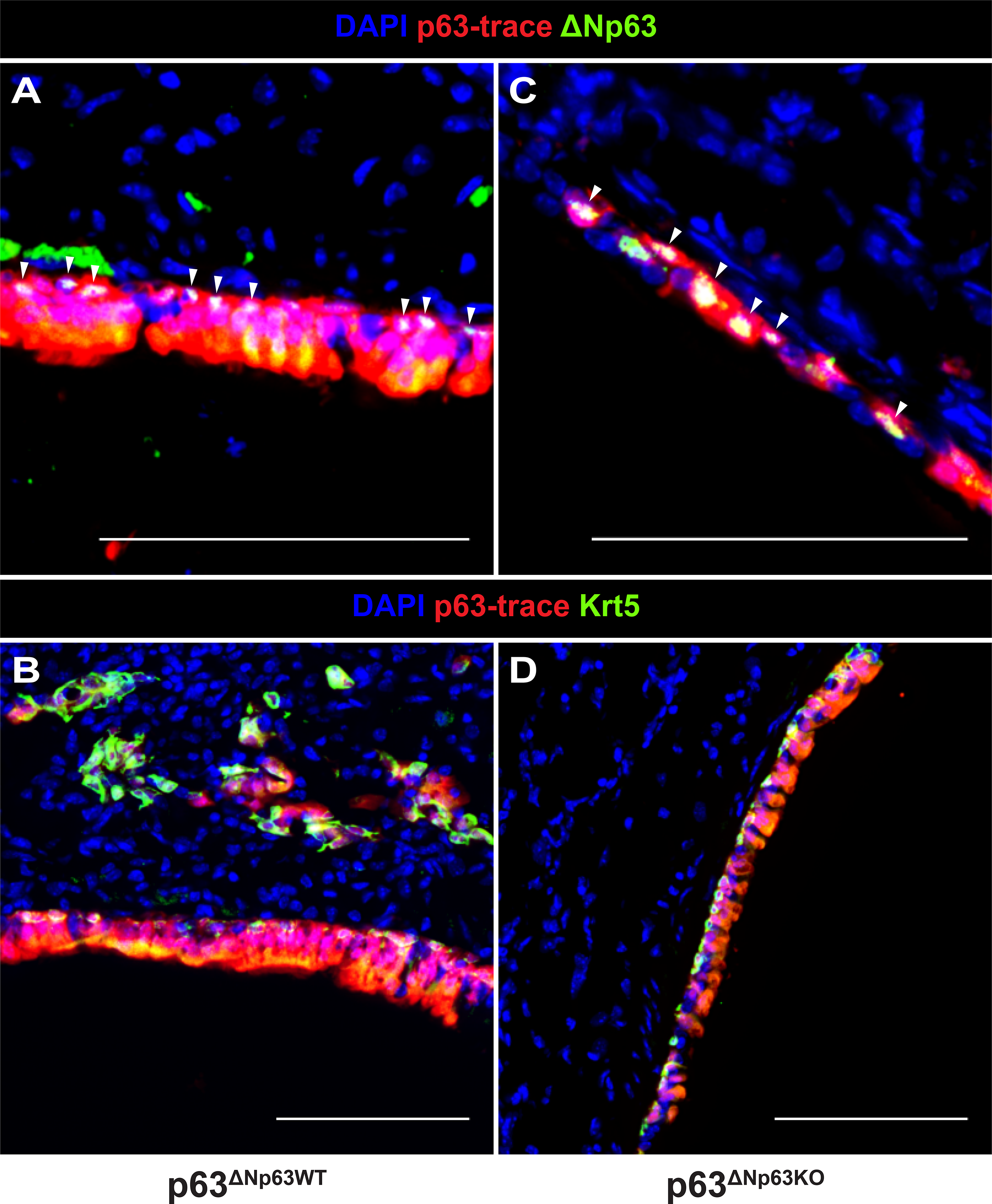
Incomplete ΔNp63^flox^ allele recombination with p63^CreERT2^ with intranasal 4OHT administration. (a, c) Immunostaining reveals ΔNp63^+^ intrapulmonary p63^+^ progenitors in the distal airways of (a) p63^ΔNp63WT^ and (c) p63^ΔNp63KO^ mice. Arrows indicate ΔNp63^+^ cells. Scale bars = 100um. (b, d) Krt5^+^ cells also remain in the distal airways of (b) p63^ΔNp63WT^ and (d) p63^ΔNp63KO^ mice. Scale bars = 100um.

**Supplementary Figure 3.**
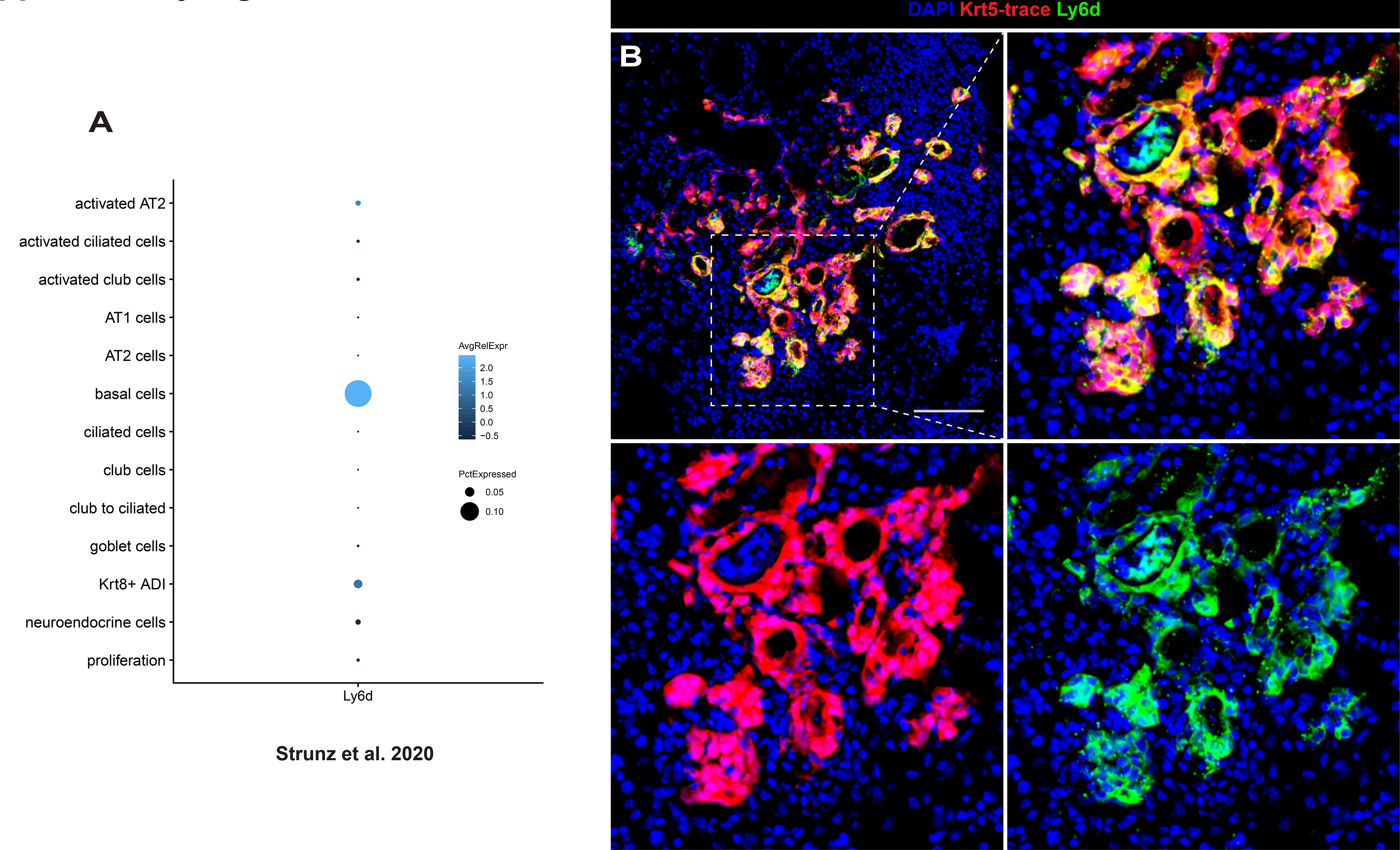
Ly6d is a cell-surface marker specific to intrapulmonary p63^+^ progenitors. (a) Dot plot from the interactive webtool (http://146.107.176.18:3838/Bleo_webtool_v2/) developed by (Strunz et al., 2020) from single-cell RNA sequencing data of bleomycin-injured murine lungs. Within the “Lung Epithelium Cell Type Signatures”, “basal cells” and “Ly6d” were queried, showing that Ly6d is highly specific to post-bleomycin intrapulmonary p63^+^ progenitors. (b) Immunofluorescence imaging of a Krt5^CreERT2^; RFP mouse given a single dose of IP tamoxifen at day 14 post-flu and sacrificed at day 21 post-flu. FITC-conjugated rat anti-mouse Ly6d staining shows high specificity of Ly7d to Krt5-trace^+^ dysplastic cells. Some Ly6d signal is also observed in nearby immune cells, which are excluded by CD45^negative^ gating in flow cytometry. Inset (right) is of outlined region. Separated channels of inset are below. Scale bar = 100um.

**Supplementary Figure 4.**
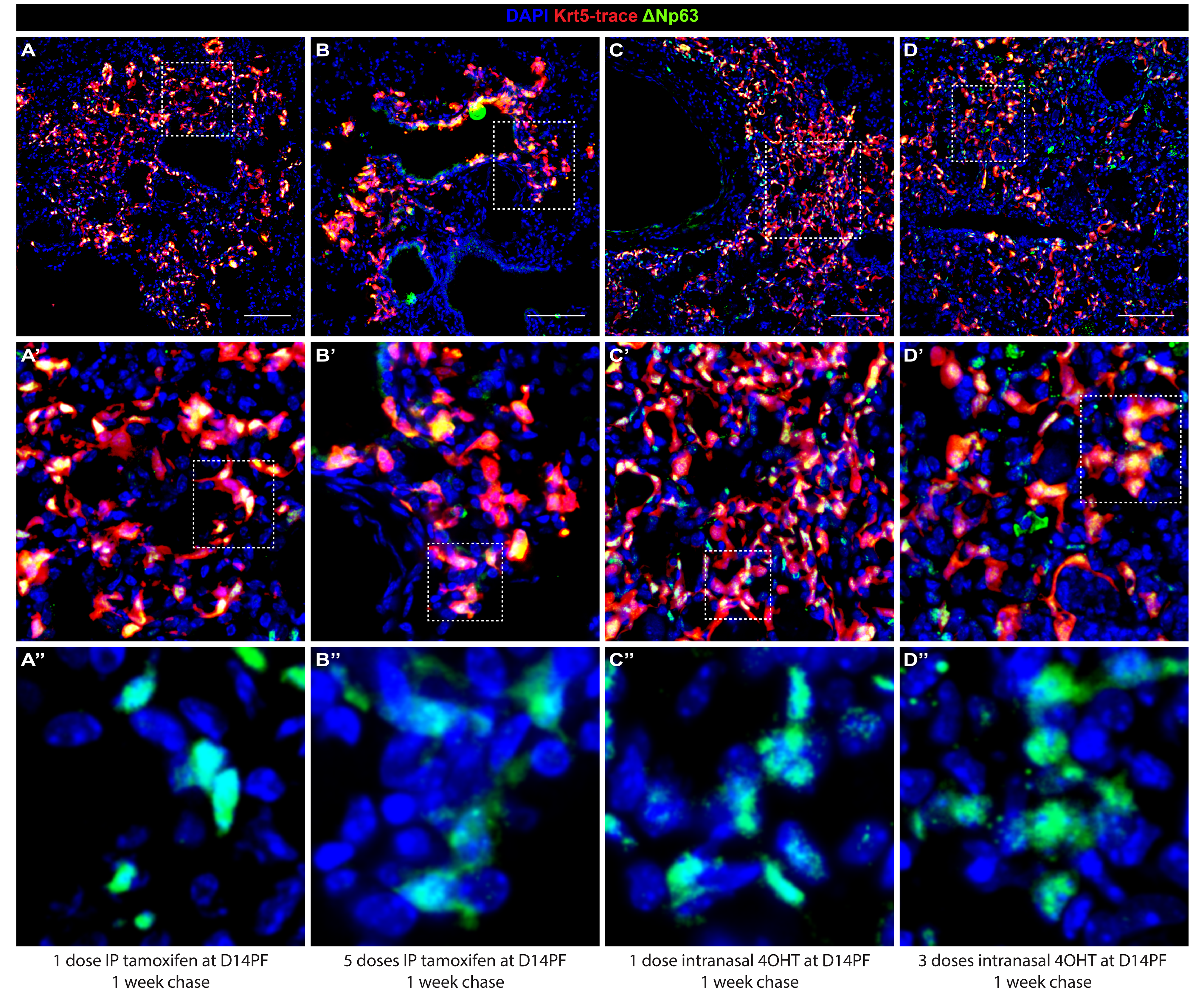
Attempts to delete ΔNp63 *in vivo*. Multiple tamoxifen dosing timelines and delivery methods were attempted using the Krt5^CreERT2^ to delete ΔNp63, including (a-a’’) one dose IP tamoxifen, (b-b’’) five doses IP tamoxifen, (c-c’’) one dose intranasal 4OHT, and (d-d’’) three doses intranasal 4OHT. In all cases, alveolar Krt5-trace^+^ ΔNp63^+^ cells were observed by immunostaining, indicating incomplete recombination of the ΔNp63^flox^ allele. Insets in (a’, b’, c’, and d’) are of outlined regions in corresponding image above. Final row (a’’, b’’, c’’, and d’’) is of indicated inset above with Krt5-trace channel removed. Scale bars = 100um.

